# Genomic imprinting of the metabolic regulator gene *Klf14* is regulated by a paternal sub-TAD anchored at *Mest* and a shared enhancer in mice

**DOI:** 10.64898/2026.05.09.724006

**Authors:** Amanda Ha, Natsuki Hayashi, Aaron B. Bogutz, Benoit Moindrot, Laura Gomez, Juliette Harris, Alexandre Marcil, Frank Court, Miyuki Shindo, Philippe Arnaud, Jacques Drouin, Thierry Forné, Daan Noordermeer, Shuji Takada, Kazuhiko Nakabayashi, Louis Lefebvre

**Affiliations:** Department of Medical Genetics, Life Sciences Institute, University of British Columbia, Vancouver, BC, Canada; Department of Maternal-Fetal Biology, National Center for Child Health and Development, Tokyo, 157-8535, Japan; Université Paris-Saclay, CEA, CNRS, Institute for Integrative Biology of the Cell (I2BC), 91198 Gif-sur-Yvette, France; Institut de Génétique Moléculaire de Montpellier (IGMM), University of Montpellier, CNRS, F-34293 Montpellier, France; Laboratoire de Génétique Moléculaire, Institut de Recherches Cliniques de Montréal (IRCM), Montréal, Québec, Canada; Genetics, Reproduction and Development Institute (iGReD), CNRS, INSERM, Université Clermont Auvergne, 63000 Clermont-Ferrand, France; Division of Laboratory Animal Resources, National Center for Child Health and Development, Tokyo, 157-8535, Japan; Department of Systems Developmental Biology, National Center for Child Health and Development, Tokyo, 157-8535, Japan; Integrated Center for Women’s Health (ICWH), National Center for Child Health and Development, Tokyo, 157-8535, Japan

## Abstract

KLF14 acts as a master regulator of gene expression in adipose tissue and variants at the human *KLF14* gene show strong and reproducible association with type 2 diabetes and metabolic syndrome. Risk alleles are only pathologic when maternally inherited, consistent with the observation that *KLF14/Klf14* is a maternally-expressed imprinted gene in human and mouse. However, how genomic imprinting is regulated at this important locus is currently unknown. In both species, *KLF14/Klf14* is located ∼200 kb away from the paternally imprinted gene *MEST/Mest*, which is regulated by a maternal gametic differentially methylated region (gDMR) at its CpG island (CGI) promoter. Although the *Klf14* CGI is kept unmethylated in most tissues, maternally-inherited DNA methylation (DNAme) marks are paradoxically required for *Klf14* expression in mouse. Here, we show that *Mest* and *Klf14* reside within the same topologically associating domain (TAD) in both mouse embryonic stem cells (ESCs) and differentiated cells, defined by sites of biallelic CTCF binding at the boundaries. Using allele-specific 4C-seq in F1 hybrid ESCs, we show that CTCF binding specifically to the unmethylated *Mest* gDMR generates a paternal allele-specific sub-TAD encompassing *Klf14*. We further show through CRISPR-Cas9-mediated mutagenesis, that deletions of the paternal *Mest* promoter region in ESCs as well as *in vivo* in mutant mice, results in both loss of *Mest* expression and acquisition of biallelic expression at *Klf14*. By analyzing epigenetic marks and chromatin looping in the *Klf14*-expressing pituitary cell line AtT-20, we identify a putative enhancer element shared by *Mest* and *Klf14*, providing a mechanistic model for the regulation of *Klf14* imprinting. This work defines a new role for the *Mest* maternally methylated gDMR, revealing that it exerts long-range effects via allele-specific modulation of TAD structures and consequently act as an imprinting control region (ICR) in the regulation of *Klf14* imprinting. Conservation of this TAD organisation at the human locus suggests that a similar regulatory mechanism operates at *KLF14*.

## Introduction

The *KLF14/Klf14* gene encodes a transcription factor of the Krüppel-like factor family^1^. Several reports have documented a strong association between variants mapping upstream of the *KLF14* gene and increased risk of developing both type 2 diabetes (T2D) and metabolic syndrome^2^. In patients of European descent, a single nucleotide polymorphism (SNP) located ∼47 kb upstream of *KLF14* is associated with susceptibility to T2D^3^. A SNP located closer to *KLF14* (∼14 kb upstream) is associated with T2D and plasma concentration of high-density lipoprotein cholesterol (HDL-C), a risk factor for coronary artery disease^4–6^. The analysis of phased Icelandic haplotypes of known parental origin identified this more proximal SNP as a T2D-associated variant correlating with lower expression of *KLF14* in adipose tissue^7^, with increased T2D risk seen only upon maternal inheritance of the C allele^8^. The strong association between the proximal SNP and expression profiles in subcutaneous adipose tissue suggested that KLF14 acts as a master *trans* regulator in this tissue^5,9^.

The manifestation of those *KLF14* T2D risk alleles only upon maternal transmission is consistent with the exclusive maternal expression of this imprinted gene in both human and mouse^10^. However, how imprinting is regulated at *KLF14* is currently unknown. In the mouse, *Klf14* is located within a large imprinted domain on chromosome 6, centered on the paternally-expressed gene *Mest* (also known as *Peg1*)^11^, and also including the *Copg2* gene, demonstrating preferential maternal expression in specific tissues^12^. The *MEST-KLF14* imprinted cluster maps to human 7q32.2 where the organization and imprinting of these two genes are conserved in humans^10,13^. Loss of *Mest* expression is associated with embryonic growth retardation and abnormal maternal behaviour, with variable penetrance depending on the mouse strain background^14–17^. Conversely, embryonic *Klf14* deficiency leads to a mild placental overgrowth with no effect on fetal growth^18^. Furthermore, both genes are associated with fat mass expansion and defects in metabolism^5,19–23^. The only known gametic differentially methylated region (gDMR) in the domain is found at the *Mest* CpG island (CGI) promoter, where a maternal CpG DNAme imprint is acquired in developing oocytes and maintained throughout development only on the maternal allele^24–26^. Maternally derived DNAme prevents transcription from this promoter specifically on the maternal allele, resulting in monoallelic imprinted expression of *Mest* exclusively from the paternal allele^11,14^. Whereas imprinted expression of *Mest/MEST* and *Klf14/KLF14* has been observed in mouse and human, the preferential maternal expression of *Copg2* has thus far only been confirmed in mouse ^10,12,27,28^. *Copg2* is adjacent and antisense to *Mest,* and imprinting of *Copg2* is observed only in the central nervous system of mice after E13.5, where *MestXL,* a longer form of *Mest,* is expressed and interferes with expression of the paternal allele of *Copg2 in cis*^29,30^. However, *MestXL* is not implicated in the regulation of *Klf14*^29^. While the mechanism underlying *Klf14* imprinting is still unknown, its expression is lost in embryos lacking global maternally inherited DNAme, paradoxically implicating oocyte-derived DNAme as essential for *Klf14* expression^10^. The *Mest* gDMR contains an allele-specific CTCF binding site, initially described in postnatal brain^31^, suggesting a potential role for this maternally methylated gDMR in structuring the genomic neighborhood of the *Mest*-*Klf14* locus in an allelic-specific manner that may be implicated in *Klf14* imprinting.

Here we investigate the chromatin structure at the *Mest-Klf14* imprinting domain in the context of sub-TAD organization. We develop an ESC *in vitro* differentiation model to study *Klf14* imprinting and apply allele-specific CTCF binding and 3D chromatin conformation assays to characterize the structure of this domain in F1 hybrid ESCs. We show that allelic CTCF binding at the *Mest* gDMR is responsible for establishing paternal allele-specific sub-TADs in the region, and subsequently that deletion of the unmethylated paternal allele leads to maternalization of the region and loss of imprinting (LOI) at *Klf14* both *in vitro* and *in vivo.* Our results establish the *Mest* gDMR as an ICR implicated in the long-range regulation of imprinted expression at *Klf14*. In support of a model implicating an enhancer shared by *Mest* (on the paternal allele) and *Klf14* (on the maternal allele), we present 3C-qPCR and Micro-C data in a pituitary cell line identifying such a candidate enhancer upstream of the *Mest* gDMR, in a region of the *Cep41* gene marked with enhancer-associated histone marks. We propose that similar allelic TAD structures and enhancer elements are responsible for the maternal expression of the *KLF14* gene in human.

## Results

### Developmental dynamics of imprinted *Klf14* expression

*Klf14* was first identified as a maternally expressed gene in human and in reciprocal C57BL/6J × JF1/Ms F1 mouse embryos^10^. *Klf14* was shown to be strictly imprinted in all mouse embryonic and extraembryonic tissues tested at E15.5, despite the lack of allele-specific epigenetic modifications at its CGI promoter at this stage. As there was no data at earlier timepoints, however, the relationship between *Klf14* imprinting acquisition and transcription in the context of development remained to be determined. To understand the regulation of *Klf14* imprinting, we mined publicly available expression datasets from F1 mouse tissues (Fig. 1a). *Klf14* shows highly dynamic expression profiles, starting weakly expressed in ESCs^32,33^, and undetectable in the inner cell mass (ICM)^33^. Post-implantation, *Klf14* expression was not detected prior to E9.5 in the embryo^34^, but visceral endoderm (VE) shows biallelic expression at this stage^32^. *Klf14* was highly expressed and imprinted in the placenta and tongue at E16.5 and P3 respectively and was also imprinted in E16.5 tissues showing lower levels of expression, such as liver and heart^32,34^. Taken together, the expression and allelic bias of *Klf14* throughout development suggest dynamic epigenetic regulation and somatic acquisition of imprinting.

**Figure 1.**
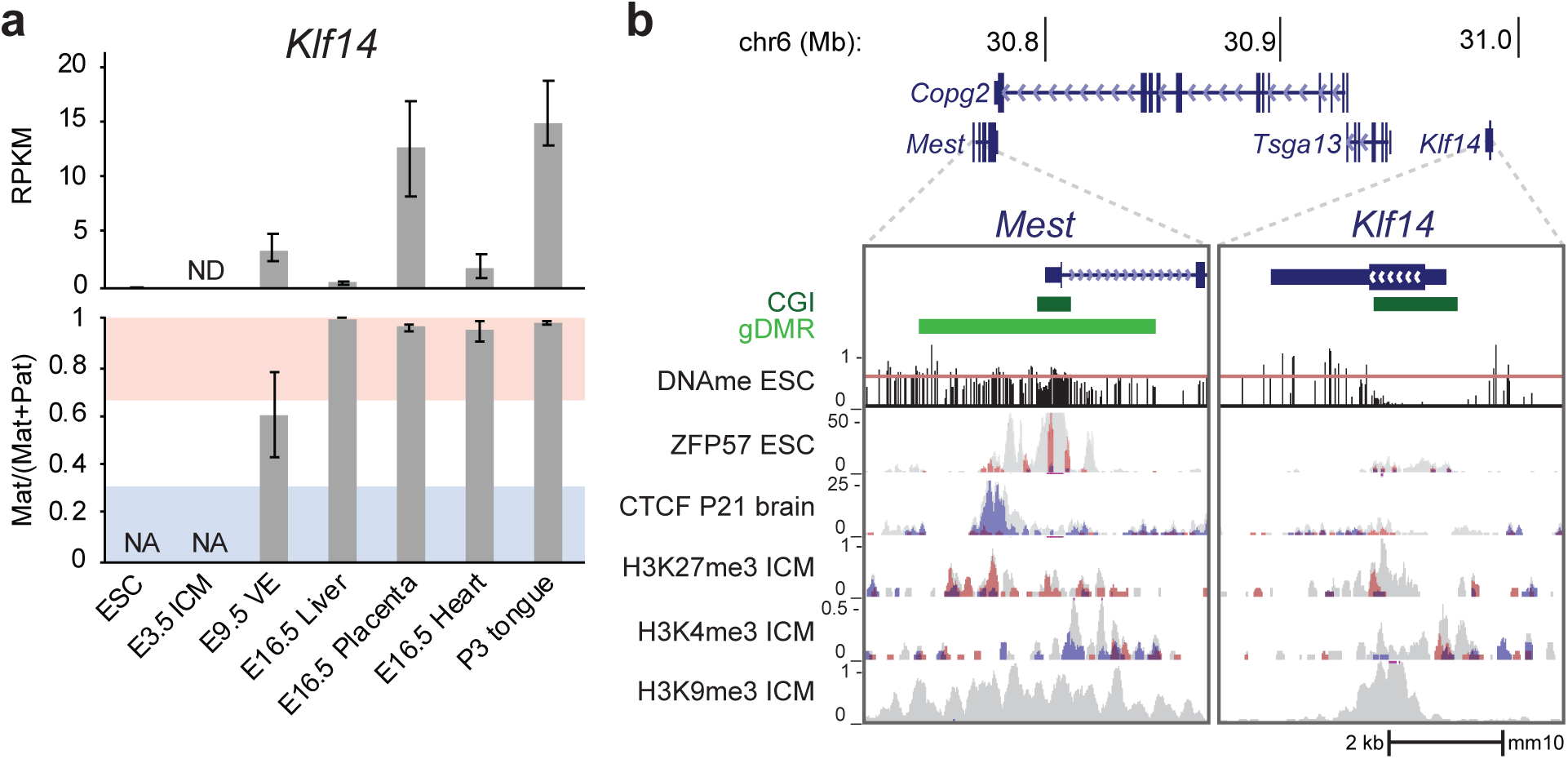
*Klf14* imprinting arises early in post-implantation development. **a**, Levels of *Klf14* mRNA and imprinting, expressed as a fraction of maternal allele expression, in mined RNA-seq data from F1 CAST × FVB mice. **b**, Visualization of allele-specific epigenetic modifications at the *Mest* and *Klf14* loci showing allele-agnostic (total; grey), and allele-specific (maternal: red, paternal: blue) modifications. Data includes DNAme, ChIP-seq for ZFP57, both in ESCs, CTCF in P21 brain, as well as histone marks: H3K27me3, H3K4me3, and H3K9me3 in preimplantation ICMs from different F1 hybrids blastocysts with C57BL/6 females. No SNP was covered for the H3K9me3 data from C57BL/6 × DBA2 mice. Details of all the datasets used in this study are presented in Supplementary Table 1. CGI: CpG island; ICM: inner cell mass.

We next investigated the allelic distribution of a variety of epigenetic marks at the *Mest* and *Klf14* promoters (Fig. 1b). As expected for an imprinted DMR of gametic origin, the *Mest* gDMR shows ∼50% DNAme in mouse embryonic stem cells (ESCs)^35^. ZFP57, a DNAme-dependent KRAB zinc finger protein responsible for maintaining allelic DNAme at gDMRs during preimplantation development^36^, is bound at the *Mest* gDMR in undifferentiated F1 ESCs specifically on the methylated maternal allele^37^. CTCF in contrast is bound specifically at the unmethylated paternal allele in adult brain^31^. In the ICM, weak expression of *Mest* is accompanied by low-level promoter marks of both H3K4me3, on the expressed paternal allele, and H3K27me3, on the maternal allele^33,38^. The gDMR is also marked by H3K9me3 in ICM, but the allelism of this mark cannot be determined due to the lack of SNPs in the mined dataset^39^.

In contrast to what is observed at *Mest*, DNAme and ZFP57 binding are absent at the *Klf14* CGI promoter in ESCs (Fig. 1b). In fact, the *Klf14* promoter remains unmethylated in all mouse tissues surveyed, as expected for a typical CGI (Supplementary Fig. 1a). Despite this, maternal DNAme was shown to be essential for *Klf14* expression in the progeny from *Dnmt3a/b*-deficient oocytes^10^. Consistent with this peculiar requirement for maternal DNAme, we find loss of *Klf14* expression in heterozygous *Dnmt3l^-/+^* E9.5 embryos (maternal allele always shown first) recovered from viable homozygous *Dnmt3l^-/-^* females, in which *de novo* DNAme fails to be established during oocyte growth (Supplementary Fig. 1b-e)^26^. No allelic H3K27me3 or H3K4me3 enrichment was found at or near the *Klf14* promoter in the ICM, consistent with previous results (Fig. 1b)^10^. No allelic information is available regarding H3K9me3 binding at the *Klf14* promoter again due to the lack of SNPs, although this mark is clearly present in ICM. However, although allelic information is not available in these datasets, the *Mest* and *Klf14* promoter regions are both marked by positive (H3K4me3) and negative (H3K9me3 and H3K27me3) histone marks, indicative of imprinting, in ESC-derived neural progenitors (Supplementary Fig. 2)^40^. Together, these observations reinforce the idea that *Klf14* imprinted expression is acquired during embryogenesis and is controlled by DNA elements distant from the *Klf14* promoter.

### Chromosome topology of the *Mest-Klf14* domain

The dependence of *Klf14* expression on maternal DNA methylation, coupled with the observations that the *Mest* gDMR harbours the only maternal allele-specific DNAme mark in this region and that it is bound by CTCF in an allele-specific manner, suggests a mechanistic role for the gDMR in the regulation of *Klf14* imprinting. To understand if the 3D organization of this genomic region may be involved in *Klf14* imprinting, we reanalyzed high-resolution Hi-C data from E14TG2a ESCs, a cell type in which TAD structures are already established (Fig. 2a)^40,41^. Our analysis revealed that *Mest* and *Klf14* are situated within the same TAD, encompassing ∼690 kb of proximal chromosome 6, from *Klhdc10* to *Klf14* (Fig. 2a). Further visual inspection revealed the presence of a weaker intra-TAD boundary at the *Mest* gDMR (Fig, 2a; arrow). Analysis of allele-specific CTCF ChIP-seq in reciprocal B6×JF1 ESCs confirmed that this intra-TAD boundary is anchored by a paternal allele-enriched CTCF peak within the *Mest* gDMR, overlapping the site previously observed in adult brain (Fig. 1b, 2a, 2b)^31^. The edges of the TAD are enriched for biallelic CTCF peaks, which may act to structure the TAD. Of note, our F1 CTCF ChIP-seq data was aligned to the mouse genome assembly GRCm38/mm10, due to the availability of SNP data, but this build contains a 50 kb genomic gap within intron 20 of *Copg2.* To address whether additional CTCF peaks map within this mm10-gap, we reanalyzed CTCF ChIP-seq from parthenogenetic (PR8) and androgenetic (AK2) ESCs^42^ and aligned these data to GRCm39/mm39, where this *Copg2* sequencing gap has been resolved. This analysis confirmed the presence of the paternal allele-specific CTCF peak at the *Mest* gDMR and a lack of additional CTCF binding within the mm10 gap (Supplementary Fig. 3). Interestingly, the CTCF binding in the promoter region of *Mest* resolves into two distinct peaks (Fig. 2b), however the upstream site is outside the gDMR, gains DNAme through development, and concomitantly loses CTCF enrichment (Supplementary Fig. 4). Together, these data show that *Mest* and *Klf14* are within the same overarching TAD and that the *Mest* gDMR harbours a paternal allele-specific CTCF binding site that may further structure the region, potentially providing differential allelic access to shared regulatory elements.

**Figure 2.**
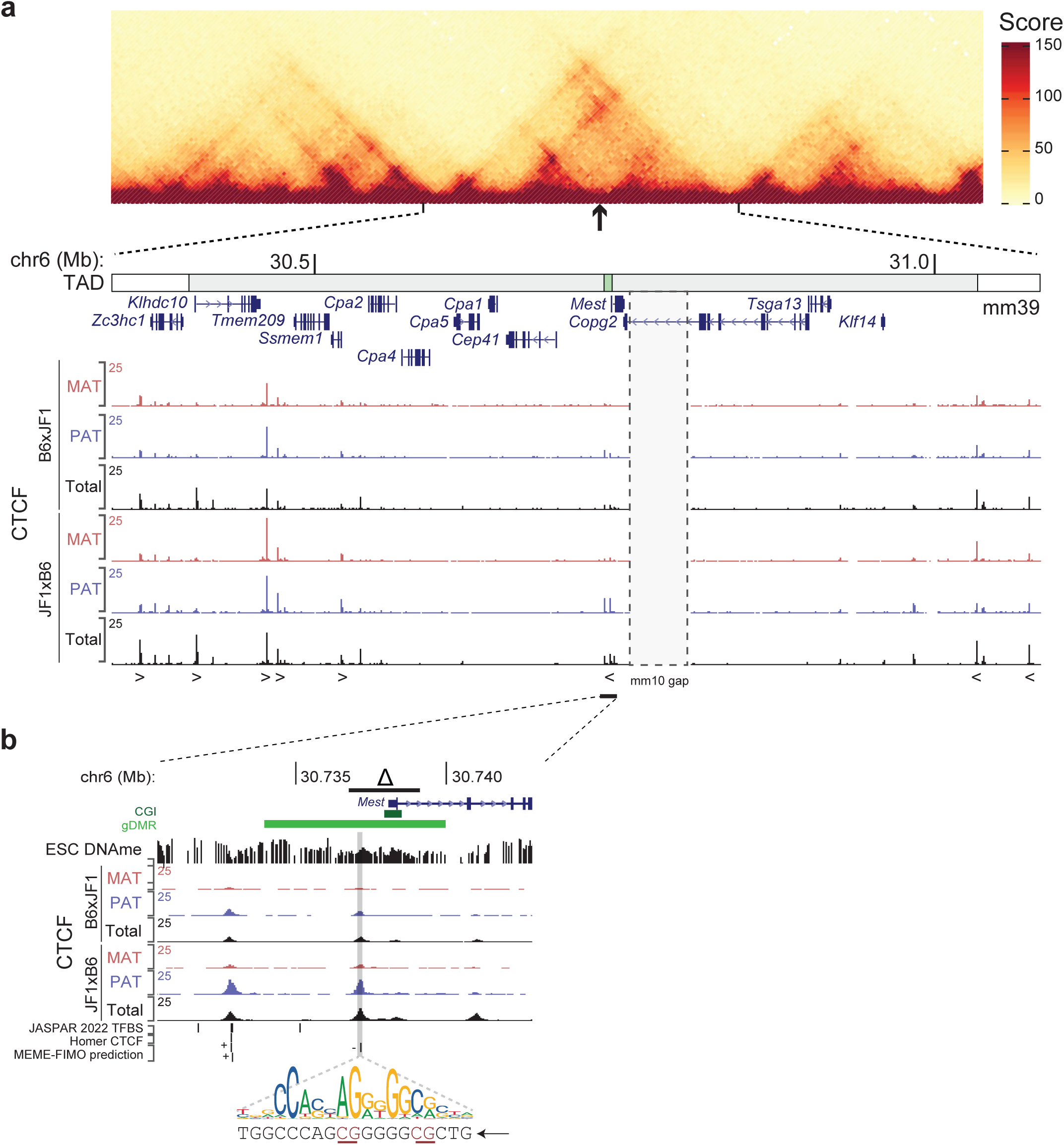
TAD organisation and CTCF binding sites in the *Mest-Klf14* region in ESCs. **a**, Hi-C signal (aligned to mm39) within the TAD encompassing *Zc3hc1*, *Mest* and *Klf14* in ESCs (grey box, indicated below the Hi-C signal), and CTCF ChIP-seq signal (aligned to mm10, which permits the distinction between the parental genomes) on the maternal (red), and paternal (blue) alleles in reciprocal B6×JF1 ESCs. The *Mest* gDMR (arrow underneath the Hi-C plot) is indicated within the TAD, above of the GENCODE gene list (green box). Note that the GRCm38/mm10 genome assembly includes a 50-kb gap within intron 20 of *Copg2*. **b**, The *Mest* gDMR encompasses a predicted CTCF binding site on the minus strand and spanning two CpGs as determined by MEME-FIMO prediction. The position of the canonical motif (grey shadow, JASPAR MA0139.1 motif) and extent of the *Mest* gDMR and CRISPR-Cas9 *Mest proKO* deletion (Δ) are highlighted above the CTCF ChIP-seq data.

### Evidence for a pituitary enhancer located upstream of the *Mest* gDMR regulating *Mest* and *Klf14*

To assess the degree of coregulation of *Mest* and *Klf14*, we analyzed data from the FANTOM5 project^43^. *Mest* expression was detected in 823 CAGE libraries while *Klf14* was detected in only 210 libraries, with *Mest* mRNA levels ranging from 10- to 10,000-fold higher than *Klf14* levels in co-expressing libraries (Supplementary Fig. 5a). Some of the highest *Klf14* levels are observed in the pituitary (Supplementary Fig. 5a, orange dots), and further analysis of *in vivo* and *in vitro* pituitary cells revealed similar *Mest* and *Klf14* expression in these cells (Supplementary Fig. 5b). To further assess the mechanism of *Mest* and *Klf14* gene regulation, we turned to the pituitary cell line AtT-20, a tissue culture model of anterior lobe corticotropes that was established from a radiation-induced pituitary tumour in a F1 hybrid C57L/J x A/J (LAF1) mouse ^44^. Analysis of published WGBS and ChIP-seq data from this cell line^45^ revealed several features indicative of imprinting at both *Klf14* and *Mest*: first, the *Mest* promoter is marked with both activating (H3K4me3 and H3K27ac) and silencing (H3K9me3) histone marks, as well as a CTCF peak in the gDMR (Supplementary Fig. 6a)^46^; second, analysis of WGBS data over the CGI promoter reveals a bimodal distribution of DNAme per sequenced reads (Supplementary Fig. 6b). WGBS reads surrounding an INDEL in intron 1 of *Mest* (absent in the parental strains) confirms allelic DNAme at this position (Supplementary Fig. 6c). A parallel analysis at *Klf14* revealed that the *Klf14* promoter is also marked with both activating (H3K4me3 and H3K27ac) and silencing (H3K9me3) histone marks (Supplementary Fig. 7a). Surprisingly, part of the promoter CGI shows a bimodal distribution of DNAme reads in WGBS data, suggesting that *Klf14* acquires a somatic DMR (sDMR) in expressing tissues (Supplementary Fig. 7b)^47^. Together, our analysis provides strong evidence that imprinting at *Mest* and *Klf14* is appropriately maintained in AtT-20 cells. Consequently, we next explored a mechanism whereby a shared pituitary enhancer, located upstream of the *Mest* gDMR CTCF boundary, might act on *Mest* on the paternal allele and on *Klf14* on the maternal allele in AtT-20 cells.

To study interaction between the *Mest* promoter and putative upstream enhancers identified in AtT-20 cells and in ENCODE data (Supplementary Fig. 5c and Supplementary Table 2), we first applied quantitative 3C-qPCR^48–50^. Using a viewpoint from the *Mest* promoter, we assayed interactions with 48 different upstream fragments, ranging from approximately 7 to 320 kb upstream of the *Mest* TSS fragment (Fig. 3a and Supplementary Table 3). Only seven fragments (dashed vertical bars), representing 3 local peaks of interactions, displayed no overlap with the local background (Fig. 3a, dashed horizontal lines). These correspond to fragments containing 4 putative enhancers numbered 6, 11, 12 and 14 (Fig. 3a, black vertical arrows, and Supplementary Fig. 5c). Utilizing high resolution Micro-C maps (Harris *et al.* and Gouhier *et al.*, manuscripts in preparation), we observed interactions between the *Klf14* and *Mest* promoter regions and the *Cep41* locus corresponding to the putative enhancers 11 and 12 identified in the 3C-qPCR analysis (Fig. 3). This region of *Cep41* is not marked by H3K4me3 (which is seen only at its promoter) but carries several hallmarks of an active enhancer, including ATAC-seq signal, H3K4me1, p300, and H3K27ac peaks (Fig. 3b and Supplementary Fig. 5c). Taken together, the 3C-qPCR and Micro-C analysis provide strong evidence for the existence of shared enhancers in the pituitary lineage acting on the *Mest* promoter on the paternal allele and the *Klf14* promoter on the maternal allele.

**Figure 3.**
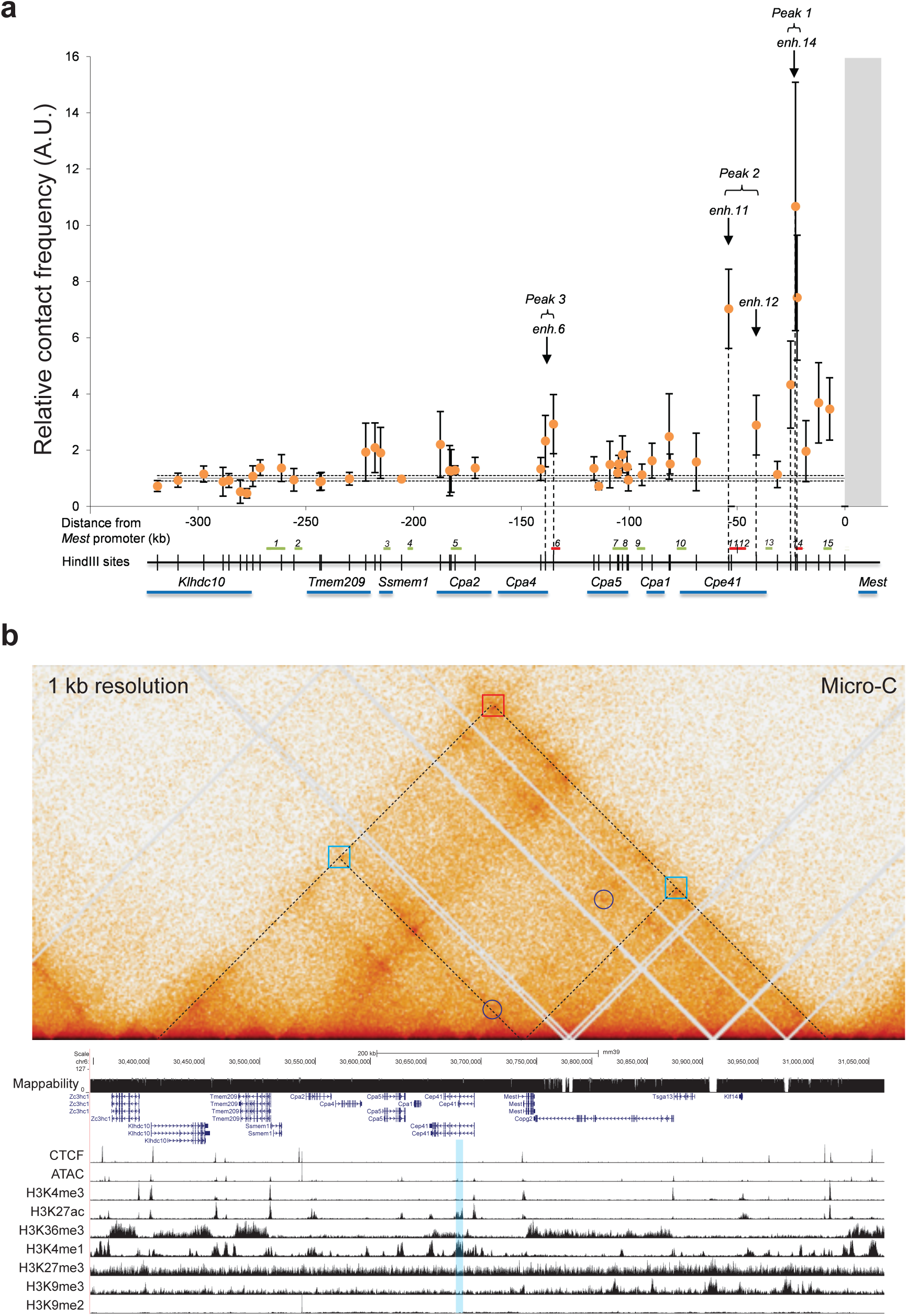
Evidence for a shared enhancer upstream of the *Mest* gDMR in AtT-20 cells. **a**, 3C-qPCR experiment showing relative contact frequencies measured between 48 HindIII fragments located upstream the *Mest* promoter (grey vertical bar), encompassing 15 putative AtT-20 enhancers (red/green horizontal bars below the graph). Dashed horizontal lines represent the local background of random contacts within the locus. Locations of HindIII sites (thin vertical bars) as well as gene positions (blue horizontal bars) are indicated below the graph. Error bars indicate s.e.m. of five independent replicates. **b,** AtT-20 Micro-C heatmap shown at 1 kb resolution. TAD structures (dashed lines) and their contact points (squares) highlight the *Klhdc10-Klf14* maternal TAD (red) and the paternal sub-TADs (blue). Interactions between the *Klf14* and *Mest* promoters with a region of *Cep41* are circled. Below: ChIP-seq data from AtT-20 cells with the putative shared enhancer region shaded in blue.

### Loss of *Klf14* imprinting upon deletion of the paternal *Mest* promoter region

For further allele-specific studies, we established a panel of reciprocal F1 C57BL/6 × CAST/EiJ ESC lines and employed early passage male ESCs with normal chromosomal counts and faithful imprinted expression of *Mest* for our studies (WT lines BC1.3 and CB1.1; Supplementary Tables 4.1, 4.2). While *Klf14* lacks detectable expression in undifferentiated ESCs, previously published microarray data reported a ∼3-fold upregulation of *Klf14* upon differentiation of ESCs to vascular progenitors (VPs) positive for receptor tyrosine kinase KDR (FLK1) expression (Supplementary Fig. 8a)^51,52^. *Flk1* embryonic expression is detected in early mesoderm and endothelial cells, coinciding with the acquisition of *Klf14* imprinting *in vivo*^53^. By RT-qPCR, we confirmed expression of *Klf14* upon differentiation of CB1.1 ESCs into FLK1^+^ VPs and detected concomitant imprinting of *Klf14,* expressed predominantly from the maternal allele (Supplementary Fig. 8b). *Mest* is also upregulated ∼6-fold in these FLK1^+^ differentiated cells^52^, suggesting that these two genes might share a similar regulatory mechanism in these cells.

To investigate directly whether the *Mest* gDMR regulates *Klf14* imprinting, we used CRISPR-Cas9 mutagenesis in our F1 ESCs to delete a ∼2.1kb region encompassing part of the *Mest* gDMR, its associated CTCF binding site and exon 1 of *Mest*, referred to below as the *Mest proKO* alleles (Fig. 2b, Δ). We produced reciprocal lines containing deletions of the paternal allele (pKO), the maternal allele (mKO), or homozygous (KO/KO) in derivatives of the BC1.3 and CB1.1 parental lines (Supplementary Fig. 9a, b). The structure of the deletions was confirmed by allele-specific analysis of each wild-type and mutant junctions (Supplementary Fig. 9c-e). *Klf14* expression was very low in undifferentiated ESCs and generally increased upon differentiation in all cell lines, although to varying degrees, likely reflecting heterogeneity of the FLK1^+^ population (Supplementary Fig. 8c). Upon differentiation of our F1 ESCs, wild-type FLK1^+^ VPs showed maternally biased *Klf14* expression (71% in BC1.3, 78% in CB1.1) (Fig. 4a). Deletion of the methylated maternal promoter has no effect on the imprinting of *Klf14*, which remains preferentially expressed from the maternal allele in FLK1^+^ VPs (Fig. 4a, mKO: 74% in CB). In contrast, deletion of the unmethylated *Mest* promoter in paternal heterozygotes (pKO) or in homozygous mutants (KO/KO) leads to loss of imprinting at *Klf14* and biallelic expression (Fig. 4a, pKO: 55.8% and KO/KO: 40.7% in BC, 51.5%, CB). These data demonstrate that *Klf14* allelic expression is dependent upon the paternal unmethylated, CTCF-bound *Mest* gDMR in differentiated VPs, while the methylated maternal gDMR is dispensable for *Klf14* imprinting.

**Figure 4.**
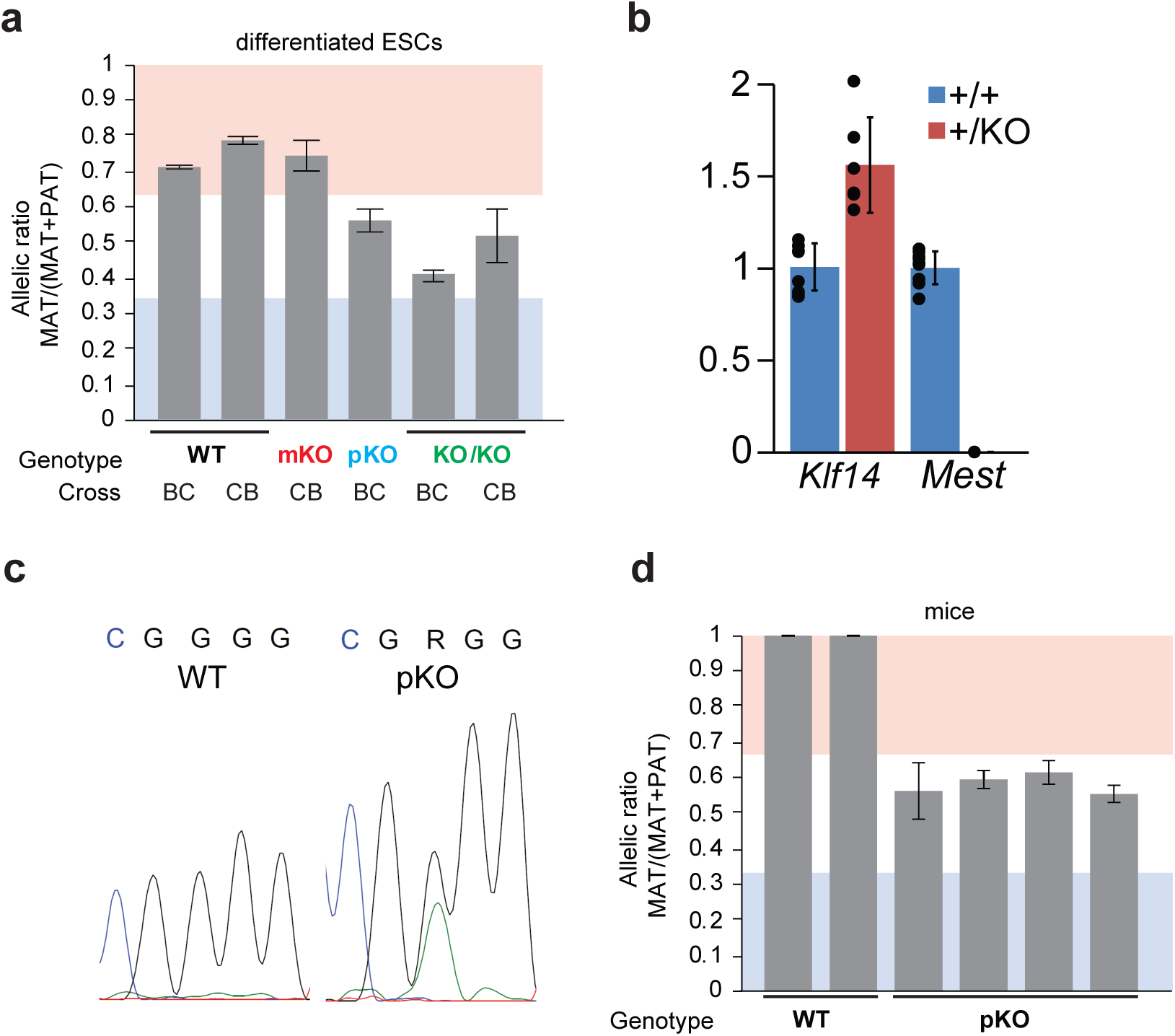
Loss of *Klf14* imprinting in *Mest^+/proKO^* ESCs and *Mest^+/gDMRKO^* embryos. **a**, Allelic expression of *Klf14* in FLK1^+^ cells from wild-type (WT) and mutant F1 ESCs. Reciprocal B6×CAST F1 ESCs (BC1.3 and CB1.1) were differentiated and sorted into FLK1^+^ populations. KO designates the CRISPR-Cas9 *Mest^proKO^* deletion alleles, encompassing the CTCF binding site and exon 1 (Supplementary Figure 9). Allelic ratios (MAT/(MAT+PAT)) of >0.65 and <0.35 represent maternal (red) and paternal (blue) expression, respectively. **b**, Expression level of *Klf14* and *Mest* in wild-type (WT) and *Mest^+/gDMRKO^* (pKO) E10.5 F1 embryos determined by RT-qPCR and expressed as a fold change from the average WT levels. **c**, Sanger sequencing trace for RT-PCR analysis of the *Klf14* SNP rs229728386 in WT and pKO F1 E10.5 embryos. **d**, Quantification of *Klf14* allelic expression ratio in individual WT and pKO embryos from a E10.5 litter.

To confirm the relationship between the *Mest* gDMR and *Klf14* imprinting *in vivo*, we generated mutant mice in which the entire ∼5-kb gDMR, as defined in gametes^54^, was deleted using CRISPR-Cas9 in zygotes (*Mest gDMRKO* alleles)^55^. Two out of approximately 60 individual mice were found to carry the expected deletion (lines 48 and 58), one of which also contains a small 947 bp ERV-rich insertion at the deletion breakpoint (line 48, Supplementary Figs. 10 and 11). Backcrosses to C57BL/6N mice generated paternal and maternal heterozygotes at the expected frequency. Maternal heterozygotes (*Mest^gDMRKO/+^)* are indistinguishable from wild-type, whereas the paternal heterozygotes (*Mest^+/gDMRKO^*) show a slight growth retardation, as has been previously described for *Mest^+/KO^* mice^14,15^. The analysis of E10.5 embryos from a wild-type JF1 female crossed to a *Mest gDMRKO* heterozygous male demonstrated that *Mest* expression is lost in the *Mest^+/gDMRKO^*mutants, as expected for imprinted expression of *Mest* and silencing of the maternal allele (Fig. 4b). *Klf14*, however, shows a modest increase in expression (Fig. 4b) accompanied by a loss of imprinting (Fig. 4c, d), suggesting upregulation of the maternal allele. The analysis of postnatal tongue, another *Klf14*-expressing tissue^32^, showed a similar pattern (Supplementary Fig. 12). Together, our results show that *Klf14* imprinting is dependent on the *Mest* gDMR both *in vitro* and *in vivo*, where it likely acts as a long-range ICR in regulating *Klf14* allelic expression.

### The *Mest* gDMR is essential for paternal sub-TAD organization

To determine the allelic nature of the TAD structures present at *Mest* and *Klf14*, we first performed allele-specific 4C-seq^56^ on reciprocal F1 ESCs using a viewpoint situated proximal to the paternal allele-specific CTCF site within the *Mest* gDMR. We observed an enrichment of paternal allele-specific interactions towards the proximal edge of a sub-TAD on the paternal allele only, with a clear switch at the *Mest* gDMR to maternal-allele specific interactions distally (Fig. 5a). Conversely, allele-specific 4C-seq using a viewpoint situated within the distal edge of the TAD, beyond *Klf14*, revealed an enrichment of interactions on the paternal allele up to the *Mest* gDMR, with another clear switch to the maternal allele proximally (Fig. 5b). Together, these observations reveal the formation of two paternal allele-specific sub-TADs within this region, hinged by the CTCF-bound paternal *Mest* gDMR (Fig. 5c).

**Figure 5.**
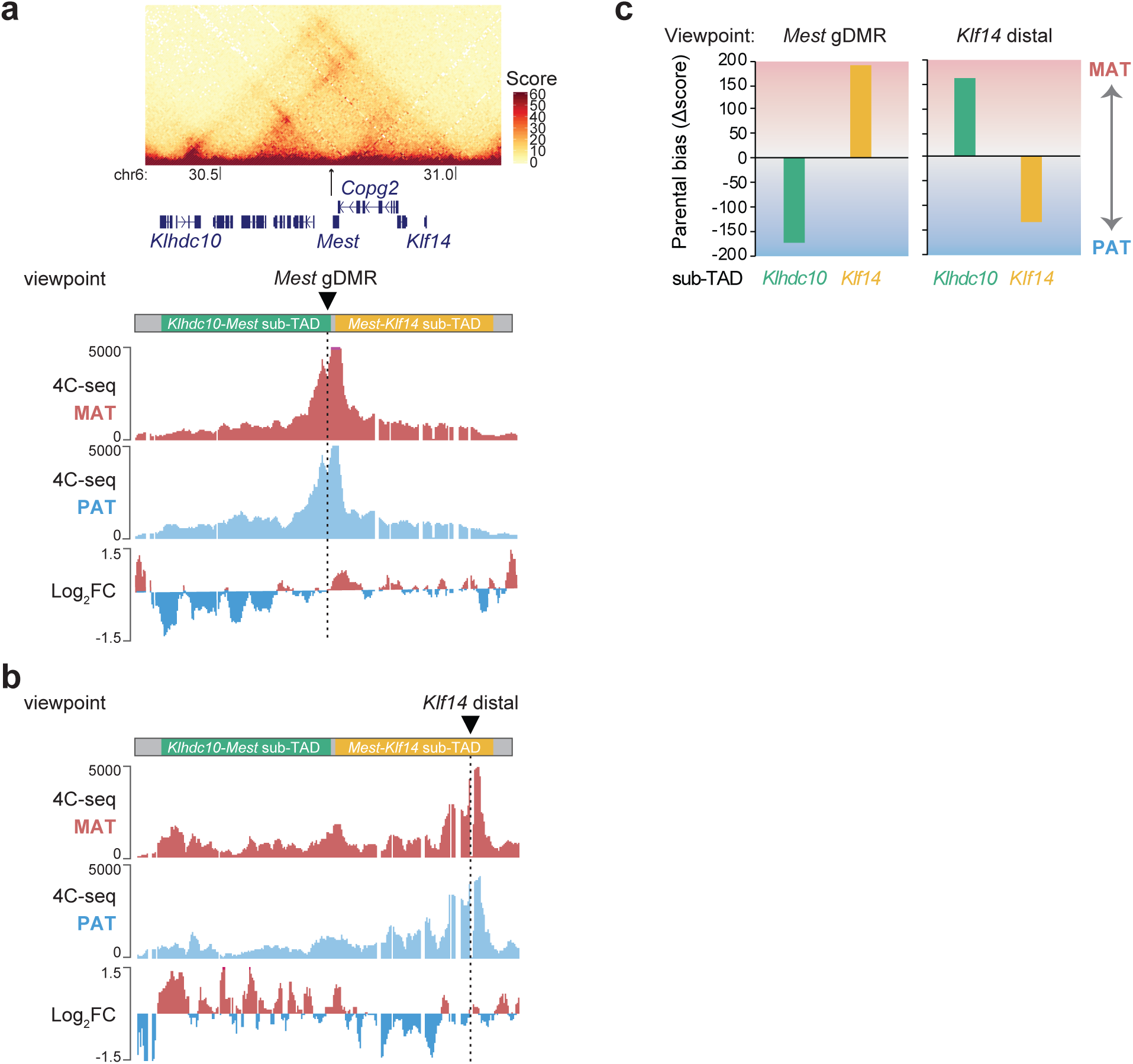
Sub-TAD organization of the *Klhdc10-Klf14* region. **a**, Allele-specific 4C-seq signal in F1 ESCs, supplemented with non-allelic Hi-C data of the *Klhdc10-Klf14* region. 4C-seq signal for the proximal *Mest* gDMR viewpoint on the maternal (red), and paternal (blue) allele in reciprocal B6×JF1 ESCs across the TAD. The log2 fold change of the above interactions is provided. The cartoon at the top depicts the sub-TAD characterized, the *Mest* gDMR viewpoint (triangle). **b**, 4C-seq signal for the distal *Klf14* viewpoint on the maternal (red), and paternal (blue) allele in reciprocal B6×CAST ESCs across the TAD. **c**, Allelic distribution of 4C-seq signal for the proximal *Mest* gDMR and distal *Klf14* viewpoints in the *Klhdc10-Mest* (347 kb) and *Mest-Klf14* (313 kb) sub-TADs. Parental bias is defined as Mat-Pat 4C-score.

To assess the role of the *Mest* gDMR CTCF site in this paternal sub-TAD, we next performed 4C-seq on the reciprocal B6×CAST ESCs with the *Mest proKO* alleles. Deletions of the promoter on the paternal allele in both pKO and KO/KO mutant ESCs strongly reduced paternal allele-specific interactions within the distal sub-TAD (Fig. 6a), while paternal interactions were increased within the gDMR-proximal *Klhdc10-Mest* sub-TAD. Conversely, there were no consistent changes in sub-TAD structure upon *Mest* gDMR deletion on the maternal allele (mKO, Fig. 6b). Quantification of the distribution of 4C-seq signal within the *Klhdc10-Mest* and *Mest-Klf14* sub-TADs revealed a loss of parental bias upon deletion of the paternal gDMR only (Fig. 6c), suggesting maternalization of the region in the pKO mutants and structural equivalence between the methylated maternal allele and the deleted paternal allele. Taken together with the presence of shared enhancers, this study provides a mechanistic model for the coupling of imprinted DNAme at the *Mest* gDMR and the allelic regulation of transcription at *Klf14* (Fig. 7), driven by allele-specific CTCF binding and resultant allele-specific TAD architecture.

**Figure 6.**
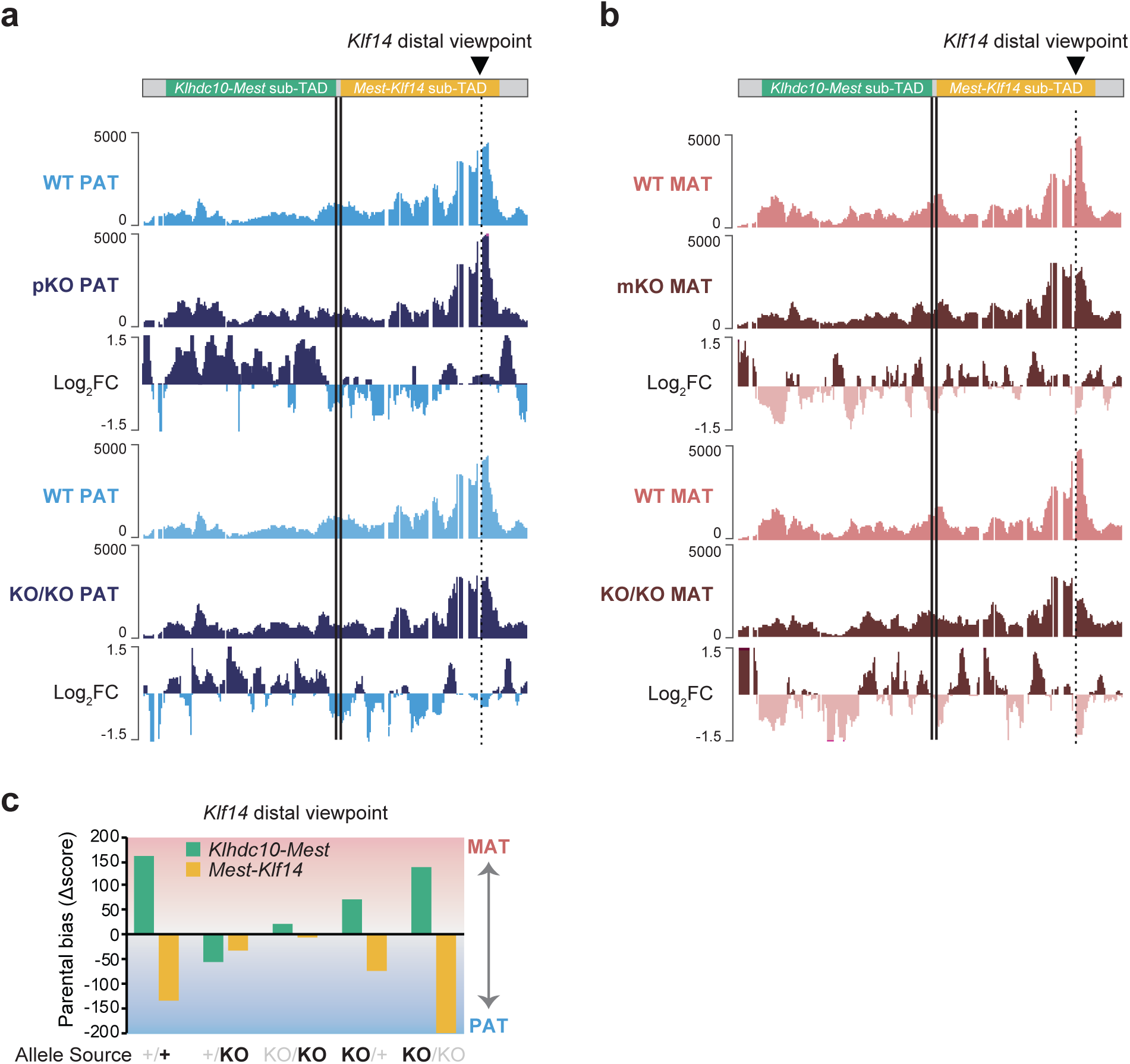
The *Mest* gDMR is essential for paternal sub-tad organization. **a**, Allelic 4C-seq signal for the *Klf14* distal viewpoint on the paternal alleles in WT and *Mest proKO* F1 ESCs. The *Mest* promoter was deleted on the paternal allele in pKO and KO/KO BC ESCs. The position of the *Mest* gDMR is highlighted by a double line. **b**, Allelic 4C-seq signal for the *Klf14* distal viewpoint on the maternal alleles in WT and *Mest proKO* F1 ESCs. The *Mest* promoter was deleted on the maternal allele in mKO and KO/KO CB ESCs. **c**, Allelic distribution of 4C-seq signal for the *Klf14* distal viewpoints in the proximal *Klhdc10-Mest* (347 kb, green) and distal *Mest-Klf14* (313 kb, gold) sub-TADs. Parental bias is defined as Mat-Pat 4C-score. The allele surveyed is in bold. The WT structures are lost in the paternal (+/**KO**, KO/**KO**), but not the maternal (**KO**/+, **KO**/KO) mutants.

**Figure 7.**
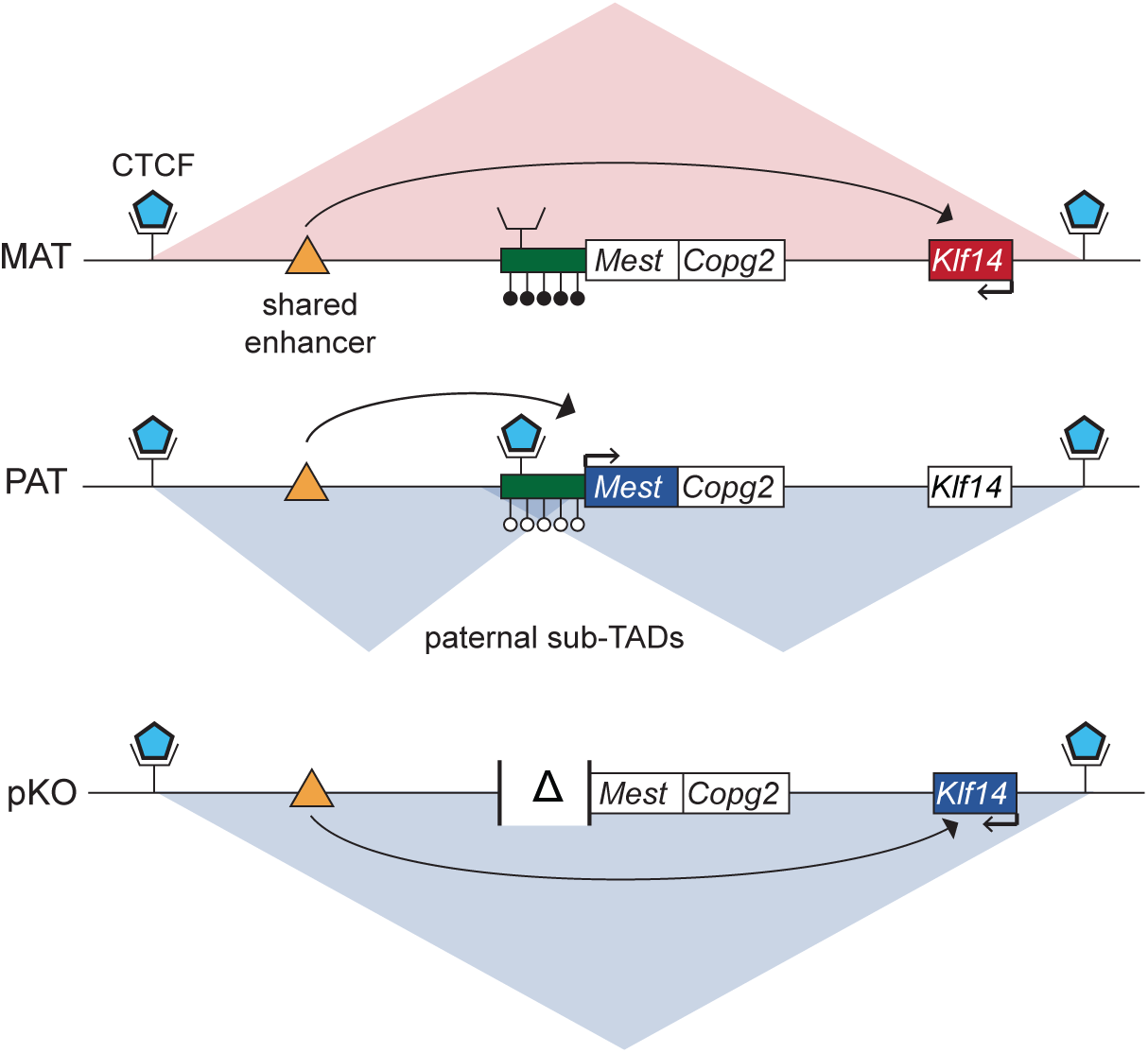
Mechanism of *Klf14* imprinting. In wild-type cells, CTCF binding at the unmethylated paternal *Mest* gDMR leads to the formation of two paternal sub-TADs, segregating *Klf14* from a shared enhancer. No sub-TAD structures are formed on the maternal allele and *Klf14* is expressed. Upon deletion of the *Mest* gDMR CTCF site on the paternal allele (pKO, Δ), *Mest* expression is abolished and sub-TAD structures are lost and *Klf14* is expressed. *Copg2* is preferentially maternally expressed only in neuronal tissues. CTCF binding (blue pentagons), DNA methylation (lollipops), putative enhancer (orange triangle) as well as maternally (red), and paternally (blue) expressed genes are indicated.

## Discussion

We report here for the first time a mechanistic model by which the *Mest* gDMR regulates imprinting of *Klf14*. CRISPR-Cas9 mutagenesis in ESCs and *in vivo* in mutant mice showed that the unmethylated allele of the *Mest* gDMR is required for both maintaining the paternal *Klf14* allele in an inactive state (Fig. 4) and establishing the paternal allele-specific sub-TADs in the region (Fig. 6). We propose that the formation of this paternal sub-TAD, defined by the differential CTCF binding at the *Mest* gDMR, segregates *Klf14* away from key putative enhancer(s) located upstream of *Mest* (Fig. 7). This model is in line with previous studies showing the importance of the structural organization of the genome via TAD formation in the establishment of regulatory neighborhoods permissive for enhancer function^57,58^. The *Mest-Klf14* imprinted domain provides a unique framework to study various aspects of monoallelic gene regulation by a single maternal DNAme imprint. The *Mest* gDMR regulates *Mest*, *Copg2*, and *Klf14* via direct promoter methylation, alternative polyadenylation^29^ and chromatin looping respectively. These observations support the hypothesis that the frequent clustering of imprinted genes is a consequence of co-regulation and highlight the convergence of multiple regulatory mechanisms at a single imprinted locus.

In our model for *Klf14* imprinting (Fig. 7), the paternal gDMR and allele-specific CTCF binding site acts as an insulator, blocking access of the *Cep41-*associated pituitary enhancer to the *Klf14* promoter. The acquisition of imprinted expression at *Klf14* might therefore correlate with the engagement of this enhancer, a model consistent with the pre-determined TAD structures, already present in ESCs, prior to the establishment of imprinted *Klf14* expression. A similar conclusion was drawn from the analysis of imprinting at the *Peg13-Kcnk9* locus^42,59,60^ and could be a general feature of CTCF-regulated imprinting^61,62^. Indeed recent reports have documented the frequent formation of sub-TADs anchored at DMRs, both gDMRs and sDMRs, in imprinted regions. The DNAme-sensitive binding of CTCF to these differentially methylated regions therefore plays an important role in mediating the long-range effects of the primary gametic imprints to several imprinted genes in the domain, via different long-range mechanisms^30,42,60,61,63,64^.

We show that paternal deletion of the *Mest* gDMR “maternalizes” the entire *Mest-Klf14* domain, suggesting a functional and structural equivalence between the maternally methylated *Mest* gDMR and a full deletion of its associated CTCF binding site. Conversely, our model suggests that loss of DNAme at the *Mest* gDMR would lead to a “paternalization” of the domain, with CTCF now binding to the maternal allele and silencing of *Klf14.* This provides an explanation for the observed loss of *Klf14* expression in embryos lacking maternally inherited DNAme genome-wide (Supplementary Fig. 1)^10^. The maternalization of the entire human domain is also implicated in the severe growth restriction Silver-Russell syndrome (SRS), with approximately 10% of patients carrying a maternal uniparental disomy for Chr7 (upd(7)mat)^65–67^. These patients are expected to be deficient for *MEST* expression and to show biallelic *KLF14* expression in expressing tissues. The mutant mice reported here with a paternally inherited *Mest gDMRKO* allele exhibit those same reciprocal effects on *Mest* and *Klf14* expression and are growth retarded. Further detailed phenotypic characterization of this mouse model will establish parallels with SRS pathologies molecularly associated with Chr7 defects, both uniparental disomy patients as well as cases with microdeletions affecting the paternal *MEST* locus^68–70^.

Although our model proposes that the paternal sub-TADs are key in keeping *Klf14* silent on the paternal allele, the relationship between TAD structure and gene expression is not always clear. For instance, it remains to be determined how *Mest* itself can be activated in the presence of those paternal sub-TADs. Recent evidence shows that promoter-proximal CTCF binding is involved in enhancer-promoter contacts, a mechanism that could be at play at *Mest*^71^. Furthermore, binding of CTCF to the *Mest* gDMR could play an important role in protecting the CGI promoter from acquiring DNAme during development, as has been observed at the maternal *H19* gDMR^72^.

In the mouse, the biallelic CTCF peak defining the distal boundary of the *Klhdc10-Klf14* maternal TAD is located ∼74 kb upstream of the *Klf14* TSS (Fig. 2a and 7), while in human ESCs, this boundary maps ∼141 kb upstream of *KLF14* (near hg38 position 130,875,500, Supplementary Fig. 13). Variants in this region of 7q32 are associated with increased susceptibility of developing cutaneous basal cell carcinoma^73^, with only minor parent-of-origin effects^8^. Based on our model, we propose that those variant-associated haplotypes interfere with the establishment of the *KLF14-*proximal TAD boundary, leading to abnormal or ectopic expression of *KLF14* as a consequence of the actions of distal enhancers in the neighbouring TAD. The disruption of CTCF-defined TAD boundaries is known to affect gene expression on either side of the deleted CTCF binding site and such a mechanism often leads to the activation of proto-oncogenes in cancer^74,75^. Consistent with a role in tumorigenesis, *Klf14* knock out mice were shown to spontaneously develop lymphomas in the spleen and lymph nodes, as well as lung adenomas^76^.

The conservation of CTCF binding and TAD structures at the *KLHDC10-KLF14* locus in human ESCs (Supplementary Fig. 13) and the detection of the paternal sub-TAD in human GM12878 cells^77^ both suggest that the model proposed here is likely to also be operative in human. Indeed, in support of our proposition of a shared enhancer, a recent analysis of the UK Biobank for parent-of-origin effects identified a *CEP41*-proximal variant (rs62471721) acting as an expression QTL on both *MEST* and *KLF14* and affecting triglyceride levels^78^. Furthermore, based on our results we propose that *KLF14* variants associated with increased risk of developing T2D upon maternal transmission are associated with haplotypes perturbing the function of the *KLF14* promoter, or its interaction with the postulated enhancer located upstream of *MEST.* Our model should provide a valuable basis for testing the relative importance of this enhancer as well as of the *MEST* gDMR and its associated CTCF boundary in regulating *KLF14* levels^79^.

## Acknowledgments

We thank Matthew Lorincz for helpful comments on the manuscript, Christine Yang for technical assistance with pyrosequencing, Tara Stach (BRC-seq, UBC) for next generation sequencing, and Justin Wong & Andy Johnson (ubcFLOW, UBC) for cell sorting. We thank Robert Feil (IGMM, Monpellier, France) for sharing of the reciprocal B6×JF1 and JF1×B6 F1 ESCs and Aurelio Balsalobre (IRCM, Montréal) for help with the AtT-20 cells. We thank Hiromi Kamura and Chiharu Tayama (NCCHD) for technical assistance with the maintenance of the *Mest* gDMR knockout mice. We acknowledge the sequencing and bioinformatics expertise of the I2BC High-throughput sequencing facility, supported by France Génomique (funded by the French National Program “Investissement d’Avenir” ANR-10-INBS-09). This work was supported by the following grants to LL: Canadian Institutes of Health Research (PJT-165992) and Natural Sciences and Engineering Research Council of Canada (RGPIN-2024-04175). Work in JD’s laboratory was supported by grants from the Canadian Institutes of Health Research (FDN-154297 and 195914) and the Digital Research Alliance of Canada (zmv-495 553). TF is supported by grants from the *ARC Fondation* (contract ARCPJA2023080006958) and the *Fondation pour la Recherche Médicale* (contract MND202310017917). This project was supported by funding from the Agence Nationale de la Recherche to DN (project IMP-Domain - ANR-22-CE12-0016) and PA (ANR-23-CE12 CORGI).

## Materials and methods

### Ethics statement

All experiments were conducted in compliance with protocols approved by the University of British Columbia (UBC) Animal Care Committee with guidelines from the national Canadian Council on Animal Care (CCAC) under certificates A19-0213 and A20-0229, and approved by the National Center for Child Health and Development (NCCHD) Animal Care and Use Committee (Permit Number: A2010-002). C57BL/6J, CAST/EiJ and JF1 mice were bred and housed in groups of two to five per cage. The mouse facility was maintained between 20-26 °C, 40-60% relative humidity, and on a 12hr light cycle.

### Analysis of *Dnmt3l* mutants

The *Dnmt3l* knockout line^80^ was maintained on a mixed 129/TerSv x C57BL/6 background. The inbred strain JF1 (Japanese Fancy Mouse 1)^81^ was obtained from the National Institute of Genetics (Mishima, Japan). Embryos at 9.5 days post coitum (dpc) were obtained from the mating of a *Dnmt3l*^+/+^ or *Dnmt3l*^-/-^ female with a JF1 male. Tail DNA was used for genotyping by PCR as previously described^80^. Total RNA was extracted from the embryos using All prep DNA/RNA mini kit (QIAGEN, 80204). RNA sequencing was performed and analyzed as previously described *(Perillous et al.,* submitted).

### ESC derivation and culture

Hybrid ESC lines BC (C57BL/6J x CAST/EiJ) and CB (CAST/EiJ x C57BL/6J) were established from F1 blastocysts as previously described^82,83^. Individual E3.5 blastocysts were isolated in M2 media, and cultured in N2B27 media with 2i (1 μM of PD0325901, 3 μM of CHIR99021) and LIF on a layer of Mitomycin-C treated mouse embryonic fibroblasts^84^. LIF was made in house from a bacterial expression vector (A. Bogutz, Lefebvre Lab). Hybrid ESC lines BJ1 (C57BL/6J x JF1) and JB1 (JF1 x C57BL/6J) were previously derived^85^ and grown as described^42^. N2B27 medium was composed of 495 mL of DMEM/F12, no phenol red (Gibco), 480 mL of Neurobasal medium (Gibco), 10 mL of B27 supplement minus vitamin A (Gibco), 5 mL of penicillin/streptomycin (Gibco), 5 mL of 200 μM L-glutamine (Gibco), 0.9 mL of 100 mM 2-mercaptoethanol, and 5mL of N2 solution. N2 solution was composed of 3.595 mL of DMEM/F12, no phenol red (Gibco), 0.5 mL of 25 μg/mL of insulin (Sigma), 0.5 mL of 100 μg/mL of apotransferrin (Sigma), 0.333 mL of BSA Fraction V (7.5%) (Gibco), 16.7 μL of 0.6 μg/mL progesterone (Sigma), 50 μL of 160 μg/mL putrescine (Sigma), and 5 μL of 3 μM of sodium selenite (Sigma). 40mL aliquots of N2B27 were stored at −80°C for up to 3 months. For all the experiments described here, we used male F1 ESC lines with normal chromosome number. ESCs were transitioned and maintained in standard ESM media (Gibco) containing 15% FBS (Hyclone, Premium Multicell), 1mM sodium pyruvate (Gibco), 0.1mM non-essential amino acids (Gibco), 0.1mM 2-mercaptoethanol (Sigma), 2mM L-Glutamine (Gibco), 50ug/mL penicillin and streptomycin (Gibco), and 0.1X LIF. All cell lines were maintained in a 37C incubator containing 5% CO2.

### Generation of *Mest proKO* F1 hybrid ESCs

Part of the *Mest* promoter gDMR, including the paternal CTCF binding site and eon 1, was deleted in reciprocal F1 ESCs using CRISPR-Cas9 to generate the *Mest proKO* alleles. We designed two single guide RNAs flanking the *Mest* gDMR CTCF binding site to generate a ∼2kb deletion: sgRNA5-1 5’-GCCCACCCCATGGCGGGATC-3’ and sgRNA3-1 5’-CTAGATCGTGCCGCGCAGTG-3’, both followed by the PAM TGG (Supplementary Fig. 9). Individual sgRNAs were designed using the Wellcome Sanger Institute Genome Editing tools (https://wge.stemcell.sanger.ac.uk/)^86^. The guides were individually cloned into a vector containing the Cas9 endonuclease and conferring puromycin resistance^87^. Vectors were transiently transfected into reciprocal F1 ESCs using Lipofectamine 3000 (Invitrogen) per the manufacturer’s instruction. Puromycin selection (1ug/mL) was started ∼12 hours later and maintained for two days. After 7-12 days of culture, individual colonies were picked, expanded, and screened for the deletion by PCR and analysed by Sanger sequencing to determine the deleted allele. PCR genotyping was performed with primers flanking the 5’ and 3’ sgRNA target sites as well as amplifying across the deletion. All primer sequences are presented in Supplementary Table 5.

### Vascular differentiation

ESCs were differentiated into FLK1^+^ vascular progenitors (VPs) as previously described^51,88^. Approximately 30,000 mESCs were plated per 35mm dish coated with collagen IV (Sigma). VP differentiation media was composed of a-MEM with no nucleosides (Sigma) with 10% FBS, 2mM L-Glutamine (Gibco), 50ug/mL penicillin and streptomycin (Gibco), and 0.05mM 2-mercaptoethanol (Sigma). Plates were incubated for 4 days with no media changes. VPs were dissociated using TrypLE Express, and incubated with 1 ug of rat anti-mouse FLK1 antibody (BD Biosciences) per 10^6^ cells for 1.5 hours at 4°C, followed by goat anti-rat Alexa488 for 30 min with propidium iodide (1X, Roche). Cells were washed with FACS buffer, and passed through a 35μm cell strainer. FACS was carried out on the Aria Cell Sorter (BDFACS) at the UBC Flow Cytometry Facility. FLK1^+^PI^-^ cells were collected. Cell pellets were collected, snap frozen in liquid nitrogen, and kept at −80 °C until use.

### Reverse transcription

Total RNA was purified using Trizol (Invitrogen Life Technologies, CA) according to the manufacturer’s directions. RNA was DNaseI-treated (Promega) at 37 °C for 1 h, and heat inactivated at 65 °C for 15 min. RNA was reverse transcribed into cDNA using M-MLV (Invitrogen) and random pentadecamers (N15).

### SNP pyrosequencing

Pyrosequencing to analyze allele-specific expression was performed using the PyroMark MD system (Qiagen) according to the manufacturer’s instructions. PCR amplification was performed using the forward biotinylated primer (Biotin-*Klf14* F3) and *Klf14 R1*. Two SNPs were analyzed to assay *Klf14* imprinted expression in BC/CB sample using amplification primers *Klf14 S1* and *Klf14 S2* (rs229728386 and rs259796680). Amplification and sequencing primer sequences are available in Supplementary Table 5. A two-tailed t-test was used to compare B6:CAST allele expression in F1 reciprocal hybrids.

### Quantitiative real-time PCR

The RT-qPCR experiments were performed in triplicate using EvaGreen dye (Biotium), and Tsg (BioBasic) on a QuantStudio 3 Real-Time PCR system (Thermo Fisher Scientific). Primers for RT-qPCR were designed using Primer Express 3.0 software, and ordered through IDT. All primer sequences are available in Supplementary Table 5. Relative expression of *Klf14* was normalized to *Ppia* expression^89^. RT-qPCR results were analyzed using LinReg PCR software using the 2^-ΔΔCT^ method^90,91^. Ct values of technical triplicates were averaged and used to calculate relative expression. As *Klf14* is an intron-less gene, RT-minus samples were run as negative controls.

### 4C-seq library preparation

Allele-specific in-nucleus 4C-seq was performed as previously described^42^ and recently detailed^56^. The MboI-NlaIII restriction enzyme pair were used. In brief, approximately 4-5 million cells were crosslinked with 2% formaldehyde for 10 minutes, and quenched with 1M glycine. Cells were washed with PBS, and the pellet was snap-frozen in liquid nitrogen for storage at −80°C. The cell pellet was lysed twice on ice for 10 min with lysis buffer: 10 mM NaCl, 10 mM Tris–HCl pH 8, 0.2% v/v NP-40 and 1X protease inhibitors. Following centrifugation for 5 minutes at 2500g (4°C), nuclei were washed with 500 uL of cold lysis buffer, and resuspended in 100 uL 0.5% SDS for 10 min at 62°C while shaking in a Thermomixer (900 rpm). Triton X-100 was added to a final concentration of 1%, and incubated for 15 min at 37°C (900 rpm). Chromatin was digested for 4 hours at 37°C (900rpm) with 400 U of MboI in 1X restriction buffer with BSA. 400 U of MboI was added every 4 hours, for a total of 1200 U incubated overnight at 37°C (900 rpm). MboI was inactivated at 65°C for 20 minutes (600 rpm). 990 uL of 1.5X ligation master mix (150 μl commercial 10X Ligation Buffer, 757 μl H2O, 75 μL 20% Triton-X100, 8 μl 20 mg/ml BSA) was added, and incubated at 16°C for 15 minutes (900 rpm). Digested chromatin was ligated using 5 uL of T4 DNA ligase (20 U/uL, Weiss units) for 4 hr at 16°C (900 rpm). Samples were de-crosslinked by incubation overnight at 65°C (900 rpm). Proteins were digested using Proteinase K and 20% SDS for 4 hr at 45°C (900 rpm). DNA was purified by phenol-chloroform-IAA extraction followed by ethanol precipitation. DNA was resuspended in 100-200 uL of ultrapure water, and incubated for 1 hour at 37°C. The second digestion was performed using NlaIII (1 U/ug of material) overnight at 37 °C (750 rpm), and heat inactivated for 20 minutes at 65°C (600 rpm). The second ligation was performed in a total volume of 14mL in 1X ligation buffer (660 mM Tris pH 7.5, 50 mM MgCl2, 50 mM DTT, 10 mM ATP) with 200U of T4 DNA ligase for 4 hours at 16 °C, followed by 30 minutes at 20–25 °C (600 rpm). DNA was purified by phenol-chloroform-IAA extraction and ethanol precipitation, and resuspended in 100-200 uL of 10 mM Tris pH 8.0, and purified using QIAquick columns (QIAgen). The 4C material was quantified by Qubit dsDNA broad range (Thermo Fisher Scientific), and stored at −20°C.

The efficiencies at each digestion and ligation step were evaluated by agarose gel electrophoresis. The resulting 4C material was amplified by inverse PCR using primers flanking the viewpoint of interest, and incorporating unique indices (Supplementary Table 3), as previously described ^92^. 4C-seq libraries were prepared using Expand Long Template PCR system (Roche) per the manufacturer’s instructions. 75 ng of DNA were used per PCR reaction, with a total of 16 individual reactions. PCR reactions were pooled, and purified using QIAquick columns.

### CTCF motif discovery

CTCF motif analysis was performed using genome-wide predictions generated by HOMER^93^ and JASPAR^94^. The CTCF *in vivo* ChIP-seq peak was obtained using the UCSC Table Browser in the mm10 genome assembly^95^. The sequence was searched for the canonical CTCF motif using the CTCF position frequency matrix obtained from JASPAR (MA0139.1) using the FIMO program of the MEME Suite^96^ and the CTCFBSDB 2.0 database^97^.

### RNA-seq and ChIP-seq analysis

Publicly available RNA-seq and ChIP-seq datasets referenced in this study are available in Supplementary Table 1. Allele-specific processing of ChIP-seq and RNA-seq was performed using MEA^98^. Processed reads were used to calculate RPKMs using VisRseq^99^. ChIP-seq was aligned using bwa^100^. Data was visualized using the UCSC Genome Browser^101^.

For CTCF allele-specific ChIP-seq in BJ1 and JB1 ESCs, published raw data^102^ were analysed as described previously^103^. Raw data for CTCF ChIP-seq in AK2 and PR8 monoparental cells are from a previous report^42^. Reads were processed as for BJ1 and JB1 cells, except that reads were mapped on the mm39 genome, in which the *Copg2* intronic gap has been resolved.

### Reanalysis of Hi-C data

Raw data were obtained from a previous publication^40^. HiC-pro^104^ was used to process the RAW reads with both the mm10 and mm39 reference genomes. Maps were generated at 5 kb resolution and normalized by the ICE-module implemented in HiC-pro and visualized using home-made scripts.

### 4C-seq analysis

The 4C-seq datasets were mapped and translated into restriction fragments using c4ctus onto mm39 (https://github.com/NoordermeerLab/c4ctus). For the *Mest* gDMR viewpoint, reads were aligned using Bowtie2^105^. For the *Klf14* distal viewpoint, reads were aligned using STAR^106^ and only uniquely mapped reads were considered. Raw 4C-seq scores were generated based on the read density across valid restriction fragments. 4C-seq scores were smoothed using a running mean across 21 MboI-NlaIII restriction fragments and then normalized based on the sum of interactions of the 5 TADs encompassing the viewpoint to allow for comparison across different samples^107^. The coordinates for TAD boundaries in mESC on mm10, called from a published dataset^40^ were converted to mm39 using LiftOver (UCSC Genome Browser). The region used for normalization on mm39 was 30.39-31.25Mb. The log2 ratio between 4C-seq scores between alleles was calculated using VisRseq^99^. BedGraphToBigWig (UCSC Genome Browser) was used to convert 4C-seq scores from bedgraph to bigwig format for visualization.

### Design and preparation of single guide RNAs (sgRNAs) for genome editing *in vivo*

We designed two single guide RNAs, sgRNA1 and sgRNA2, to generate a deletion corresponding to the *Mest* germline differentially methylated region (chr6:30734933-30739966 (mm10)) as previously defined at the *Mest* promoter.^54^. The target genomic sequences of sgRNA1 and sgRNA2 are 5’-GAGCAAAAAAGCACAGGATT<u>AGG</u>-3’ (mm10, chr6:30734970-30734992) and 5’-GTGACCATGGCGGGTCACAA<u>AGG</u>-3’ (mm10, chr6:30739914-30739936), respectively. These target genomic DNA sequences contain the PAM sequence (AGG) at their 3’ end (underlined). The sgRNAs were prepared as described previously by Inui *et al.*^55^. The primer sequences used for sgRNA cloning were sgRNA1-F and sgRNA1-R for sgRNA1 and sgRNA2-F and sgRNA2-R for sgRNA2. The primers to obtain PCR-amplified templates for *in vitro* transcription were gRNA1-F for sgRNA1, gRNA2-F for sgRNA2, and gRNA-R as a common reverse primer for both sgRNAs. Human codon-optimized Cas9 (hCas9) and sgRNA cloning vectors were gifts from George Church (Addgene Plasmids #41815 and #41824, respectively). The hCas9 mRNA was prepared as described previously^55^. Briefly, *in vitro* transcription was performed using the mMESSAGE mMACHINE T7 Transcription Kit (Invitrogen, AM1344) with PCR-amplified templates generated from the hCas9 plasmid using the primers T7 Cas9 Fw and Cas9 Rv. The resultant hCas9 mRNA was purified using the MEGAclear Transcription Clean-Up Kit (Invitrogen, AM1908). Primer sequences are available in Supplementary Table 5.

### Microinjection

The microinjection of gRNA1, gRNA2 and Cas9 mRNA into mouse zygotes was performed as described previously^55^. The two sgRNAs and Cas9 mRNA were mixed at 1:1:1 ratio (167 ng/μl each), and injected to the mouse zygotes obtained by mating superovulated BDF1 females and WT BDF1 males (Sankyo lab service). The protocols for animal experiments were approved by the Animal Care and Use Committee of the National Center for Child Health and Development (A2009-002-C04).

### Nucleic acid extraction

Genomic DNA was extracted from mouse tails using Quick-DNA 96 Kit (ZymoResearch, D3012). Total RNA was extracted from mouse embryos and tissues using AllPrep DNA/RNA Mini Kit (Qiagen, 80204) or AllPrep DNA/RNA Micro Kit (Qiagen, 80284).

### PCR genotyping of *Mest gDMRKO* mice

The genotyping of individual mouse of *Mest*-DMR knockout lines was conducted by PCR using primers KN156 and KN157, which amplify a 5626 bp fragment from the wild-type *Mest* allele (mm39, chr6:30,734,632-30,740,257). This 5626 bp amplicon contains the sgRNA1 and sgRNA2 sequences. PCR amplification was performed on genomic DNA from mouse tails in 20 µL reactions using PrimeSTAR® HS DNA Polymerase (Takara, R010A) with cycling conditions of: 98°C denaturation for 20 s; 30 cycles of 10 s at 98 °C, 15 s at 63 °C, and 1 min at 68 °C. Primer sequences are available in Supplementary Table 5. The knockout lines were backcrossed onto the C57BL/6N background for more than 10 generations prior to analysis.

### Allelic expression analysis by Sanger sequencing

Allelic expression analysis was conducted for tissues from F1 hybrid embryos or post-natal individuals obtained by the cross of a wild-type JF1/Ms female and a male of *Mest* DMR knockout line heterozygous for the deletion. JF1/Ms was obtained from the National Institute of Genetics (Mishima, Japan). SNPs rs229728386 at chr6:30957834 (mm10) and rs264739018 at chr6:30745854 (mm10) were used to distinguish parental alleles of *Klf14* and *Mest*, respectively. The nucleotide bases of the SNPs on the sense strand of the reference genome sequence are T/C (rs229728386) and G/A (chr6:30745854) for the C57BL6 and the JF1 strains. cDNA amplicons containing the SNP sites were amplified for *Klf14* and *Mest* using primers KN250/KN251 and KN252/KN253, respectively, with Ex Taq® Hot Start Version (Takara, RR006A). After cleanup using ExoSAP-IT or AMPureBeads, the PCR products were subjected to sequencing reactions using BigDye Terminator v3.1 Cycle Sequencing Kit, followed by purification using BigDye XTerminator Purification kit. Sanger sequencing was conducted on a ABI3500xl platform. Primer sequences are available in Supplementary Table 5. To measure *Klf14* allelic ratios, Phred (http://www.phrap.org/phredphrapconsed.html) was run on .ab1 files using the “-d” option to output polymorphism data. Quantification was performed using the calculated relative areas of the called and uncalled bases at SNP sites^108^. Data shows average and standard deviation calculated from forward and reverse sequencing technical replicates.

### Reverse-transcription quantitative PCR (RT-qPCR)

Total RNAs were extracted from the F1 hybrid embryos or post-natal tissues of F1 hybrid individuals obtained by the cross of a wild-type JF1/Ms female and a male of *Mest*-DMR knockout line heterozygous for the deletion. JF1/Ms was obtained from the National Institute of Genetics (Mishima, Japan)^81^. One microgram of total RNA was subjected to first-strand cDNA synthesis using PrimeScript™ RT reagent Kit with gDNA Eraser (Takara, RR047A). Real-time PCR reactions were performed with technical duplicates on the 7500 Fast Real-Time PCR System (Applied Biosystems, Foster City, CA, USA) using TB Green® Premix Ex Taq™ II (Tli RNaseH Plus) (Takara, RR820A) according to the manufacturer’s instructions. PCR primers were as follows: KN250 and KN251 for the first primer pair targeting *Klf14*; KN341 and KN342 for the second *Klf14* primer pair; KN252 and KN253 for *Mest*; and Actb-F and Actb-R for *Actb*. For each reaction, one-fiftieth of the synthesized cDNA was used in a 20 µL reaction volume. Real-time PCR was performed under the following conditions. For *Mest* and *Actb*, amplification was carried out with an initial denaturation at 95°C for 20 s, followed by 40 cycles of 95°C for 3 s and 60°C for 30 s (two-step cycling). For *Klf14*, amplification was performed with an initial denaturation at 95°C for 20 s, followed by 40 cycles of 95°C for 3 s, 64°C for 30 s, and 72°C for 30 s (three-step cycling). A melt curve analysis was performed after amplification to confirm product specificity by increasing the temperature from 60°C to 95°C with continuous fluorescence acquisition. Relative expression levels of *Klf14* and *Mest* were calculated by the ΔΔCt method using the Ct values of *Actb* as a reference gene. Primer sequences are available in Supplementary Table 5.

### Flongle sequencing of PCR amplicons

PCR amplicons were purified and quantified prior to library preparation. Sequencing libraries were prepared using the Ligation Sequencing Kit (SQK-LSK114, Oxford Nanopore Technologies) in combination with the Native Barcoding Expansion 1–12 (EXP-NBD104), according to the manufacturer’s instructions. Barcoded libraries were pooled and sequenced on a Flongle flow cell (FLO-FLG114) mounted on a GridION device.

### Amplicon sequencing to determine allelic ratios

PCR amplicons (1 ng input DNA) were purified using AMPure XP beads (Beckman Coulter, A63881) according to the manufacturer’s instructions. Sequencing libraries were prepared using the NEBNext Ultra II DNA Library Prep Kit for Illumina (New England Biolabs, E7645S) following the manufacturer’s protocol. Briefly, purified PCR products were subjected to end repair and adaptor ligation, followed by library amplification with 6 cycles of PCR. The resulting libraries were purified using AMPure XP beads, and sequenced on an Illumina MiSeq platform using the MiSeq Reagent Kit v2 nano (Illumina, MS-103-1001) according to the manufacturer’s instructions.

### 3C-qPCR analysis of AtT-20 cells

The 3C-qPCR experiment and data normalization have been performed as described previously^109^, including controls for digestion efficiencies, sample purity, and efficiencies of qPCR primers. The BACs RP24-275J18 and RP24-286P12 were used to generate a control template matrix, following HindIII digestion and ligation ^110^. Experiments have been performed on 5 samples each containing 5 million nuclei prepared from ATt-20 cells. Using a reverse bait primer upstream of the *Mest* promoter (mm39, chr6:30,731,554-30,731,575), and immediately downstream of the last HindIII site before the *Mest* CGI promoter, we tested 48 upstream HindIII fragments for promoter interactions. The exact values of interaction frequencies and error bars for each reaction, as well as the sequences of all 3C-qPCR primers used in this experiment are provided in Supplementary Table 3. The Noise Band of the experiment (horizontal dashed lines in the Fig. 3) is: 1 +/- 0.096.

### AtT-20 data

All WGBS and ChIP-seq data from AtT-20 cells have been published^45,111,112^. The relevant datasets (Supplementary Table 1) were aligned to mm10 or mm39 using a custom script to generate BAM files for analysis and BigWig files for visualisation in the UCSC Genome Browser^113^. BAM files from the WGBS data were intersected to specific regions of the *Mest* and *Klf14* loci using BEDTools^114^ and visualised with IGV^115^. Generation of the Micro-C data in the laboratory of JD will be published elsewhere (Harris *et al.* and Gouhier *et al.*, manuscripts in preparation).

---

**Supplementary Figure 1. *Klf14* expression requires a maternal DNAme mark. a**, Absence of DNAme at the *Klf14* promoter CGI. DNAme profiles at the *Klf14* locus, showing WGBS data from mouse germ cells and adult tissues. Blue lines highlight annotated hypomethylated regions. **b**, Genome Browser view of *Klf14* showing allelic RNA-seq signals in representative wild-type (WT) and *Dnmt3l^-/+^* E9.5 embryos. Total and merged parental allelic signals are displayed in the top and bottom panels, respectively, with maternal and paternal expression levels shown in red and blue, respectively. **c**, *Klf14* expression levels, measured as RPKM from RNA-seq, in WT E9.5 (n = 7) and *Dnmt3l^-/+^* (n = 6) E9.5 embryos from homozygous females. **d**, Ratio of maternal *Klf14* expression in WT (n = 7) and *Dnmt3l^-/+^* (n = 6) E9.5 embryos, calculated as the fraction of RNA-seq reads derived from the maternal *Klf14* allele. Insufficient signal at informative allelic sites was detected in *Dnmt3l^-/+^* embryos to establish an allelic ratio. **e**, Heatmap showing the expression Z score (upper panel) and allelic levels (lower panel) of *Klf14* in E9.5 WT (left panel) and *Dnmt3l^-/+^* embryos. Data are presented for each independent embryo (WT: n=7; *Dnmt3l^-/+^*: n=6). Insufficient signal at informative allelic sites was detected in *Dnmt3l^-/+^*embryos to establish an allelic ratio.

**Supplementary Figure 2. Epigenetic marks at *Mest* and *Klf14* in NPCs.** Histone modifications centered on the *Mest* (**a**) and *Klf14* (**b**) promoter CGIs, in ESC-derived NPCs. Details of all the datasets used in this study are presented in Supplementary Table 1. CGI: CpG island. NPC: neural progenitor cell.

**Supplementary Figure 3. CTCF ChIP-seq from monoparental parthenogenetic and androgenetic ESCs.** CTCF ChIP-seq data from undifferentiated monoparental ESCs, aligned to the mm39 mouse genome build. Position of the sequencing gap present in mm10 is highlighted in blue at the top. The bottom panel shows a magnified view of the *Mest* promoter region and exon 1 CGI. PR8: parthenogenetic; AK2: androgenetic. Details of all the datasets used in this study are presented in Supplementary Table 1.

**Supplementary Figure 4. Analysis of allelic CTCF binding sites near the *Mest* CGI promoter. a**, UCSC Genome Browser screenshot of DNAme, and CTCF ChIP-seq in F1 ESCs and somatic tissues. Predicted CTCF binding sites from HOMER and JASPAR are indicated below. The canonical CTCF binding motif (JASPAR MA0139.1) predicted by MEME-FIMO is highlighted over the upstream (u1, u2) and gDMR (g1) sites. **b**, MEME-FIMO motif analysis of the predicted CTCF binding sites in the upstream peak. The canonical CTCF motif on the (+) strand is indicated and overlapping CpG sites are underlined. **c**, Quantification of mean DNAme over the upstream and *Mest* gDMR CTCF binding sites in ESCs and somatic tissues (heart, lung). Details of all the datasets used in this study are presented in Supplementary Table 1.

**Supplementary Figure 5. Expression and epigenetic data in pituitary cell types. a**, FANTOM5 data on *Mest* and *Klf14* expression, expressed as CAGE library normalized counts, in 210 different libraries in which *Klf14* is expressed. Data from the pituitary, from embryos and neonates, are highlighted in orange. Axes also show the density plots. **b**, Expression of genes from the *Klhdc10-Klf14* TAD in purified corticotropes and AtT-20 cells. **c**, UCSC Genome Browser screenshot from mm10, highlighting epigenetic data from AtT-20 corticotrope-like cells, showing the positions of enhancers associated with the *Mest* promoter in the ENCODE 3 data as well as a set of 15 putative AtT-20 enhancers located upstream of the *Mest* gDMR.

**Supplementary Figure 6. Epigenetic marks at *Mest* in AtT-20 cells. a**, UCSC Genome Browser screenshot showing epigenetic marks at the *Mest* promoter region in AtT-20 cells, including the positions of the *Mest* CGI and gDMR as defined in E12.5 embryos. **b**, AtT-20 WGBS reads interacting with the Mest CGI shown in IGV, with CpG sites coloured (red, methylated; blue, unmethylated). Inset shows violin plot of the distribution of mean DNAme levels per strand for those 145 reads. **c**, AtT-20 WGBS reads viewed as pairs in IGV and coloured to highlight those with (orange) or without (pale blue) a heterozygous 1-bp insertion within intron 1 of *Mest*, detected in AtT-20 cells.

**Supplementary Figure 7. Epigenetic marks at *Klf14* in AtT-20 cells. a**, UCSC Genome Browser screenshot showing epigenetic marks at the *Klf14* promoter region in AtT-20 cells. **b**, AtT-20 WGBS reads interacting with a portion of the *Klf14* CGI shown in IGV, with CpG sites coloured (red, methylated; blue, unmethylated). Inset shows violin plot of the distribution of mean DNAme levels per strand for those 110 reads.

**Supplementary Figure 8. Expression and imprinting of *Klf14* in ESC-derived FLK1^+^ vascular progenitors. a**, Schematic of *in vitro* differentiation of CB1.1 ESCs into mesoderm and flow cytometry data representation. The proportion of FLK1^+^ vascular progenitors sorted is indicated. **b**, Imprinting and expression levels of *Klf14* during *in vitro* differentiation of F1 CAST female × B6 male (CB1.1) ESCs into FLK1^+^ vascular progenitors. Error bars represent the SD of technical triplicates. ND: not detected. **c**, Characterization of *Klf14* expression levels upon deletion of the *Mest* gDMR CTCF site *(Mest^proKO^* allele) in reciprocal B6×CAST ESCs differentiated into FLK1^+^ vascular progenitors, by qRT-PCR normalized to *Ppia*. Results for sorted FLK1^+^ populations are shown. Error bars represent the SD of qPCR triplicates. B6×CAST and CAST×B6 F1 cell lines with C57BL/6 and CAST maternal genomes, respectively. The CRISPR-Cas9 *Mest^proKO^* deletion alleles (KO) are as presented in Supplementary Figure 9. Maternal (KO/+) and paternal (+/KO) deletions are labeled as mKO and pKO, respectively.

**Supplementary Figure 9. CRISPR-Cas9 deletions of the *Mest* promoter. a**, UCSC Genome Browser tracks showing the 5’ end of the somatic *Mest* transcripts, the CGI associated with exon 1 (green), the position of the paternal CTCF peak (blue) and the extend of the CRISPR-Cas9 deletions characterized in four different ESC lines, derived from the BC1.3 and CB1.1 parental lines. The positions of the two gRNAs are shown below (red). **b**, DNA sequence of each promoter deletion allele *Mest^proKO^* (or KO for short), highlighting the CRISPR sgRNAs in red, the PAM sequences underlined and in bold, and a 3-bp insertion in CB1.1 KO/KO in lowercase letters. Maternal (KO/+) and paternal (+/KO) deletions are labeled as mKO and pKO, respectively. The two homozygous lines have the same deletion on both alleles. Dotted line: intervening nucleotides in WT DNA sequences. Solid line: deleted sequences (not to scale). See Material and Methods for allele nomenclature, deletion sizes, and mm39 coordinates. **c**, Design of three PCR assays to analyse ESC genomic DNA of *Mest proKO* clones. **d**, PCR and sequencing primers for each reaction. **e**, Informative parental BC SNPs monitored. All primer sequences are in Supplementary Table 5.

**Supplementary Figure 10. Structure of the *Mest gDMRKO* allele. a**, UCSC Genome Browser window showing the structure of the *Mest gDMRKO* allele, a 4940-bp deletion encompassing the paternal gDMR CTCF binding site as well as the *Mest* exon 1 CGI. The ESC *Mest proKO* allele is shown for comparison. Bottom tracks show CTCF ChIP-seq data from BJ1 ESCs, with tracks for the maternal B6 allele (MAT) and paternal JF1 allele (PAT) shown separately. **b**, Structure of the 947-bp ERV-rich insertion at the deletion breakpoint in the mutant mouse line 48.

**Supplementary Figure 11. *Mest gDMRKO* allele in mouse line 48.** DNA sequence analysis of the ∼1.6 kb fragment amplified from genomic DNA of *Mest gDMRKO* mouse line #48, using PCR primers KN156/157 (see Supplementary Table 5).

**Supplementary Figure 12. Loss of imprinting at *Klf14 in vivo* in P2 tongue. a**, PCR genotyping of P2 tongue samples from two litters obtained from crosses between a JF1 female and male *Mest^+/gDMRKO^* line #48. **b**, Sanger sequencing traces around the *Klf14* SNP amplified by RT-PCR with primers KN250 and KN251 on P2 tongue samples. **c**, Allelic ratios (Mat/(Mat+Pat)) in RT-PCR products determined by Phred analysis of Sanger reads. **d**, Allelic composition at the *Klf14* SNP in RT-PCR products from F1 E10.5 embryos and P2 tongue samples, determine by Flongle sequencing of amplicons. **e**, Fold change in levels of *Klf14* and *Mest* mRNAs, amplified by RT-qPCR with the primers indicated (top) in WT (+/+) and *Mest^+/gDMRKO^*(+/KO) P2 tongue samples.

**Supplementary Figure 13. TAD structure of the *KLHDC10-KLF14* region on human chromosome 7q32 in hESCs.** ChIP-seq data for CTCF and for the cohesin complex component RAD21 as well as Micro-C data for the *MEST-KLF14* region in undifferentiated H1 hESCs, aligned to human genome build GRCh38. CTCF and RAD21 sites defining the boundaries of the *KLHDC10-KLF14* TAD and sub-TADs are highlighted in green and blue, respectively.

