## Supplement for "Genomic imprinting of the metabolic regulator gene *Klf14* is regulated by a paternal sub-TAD anchored at *Mest* and a shared enhancer in mice"

### Supplementary Figure 1

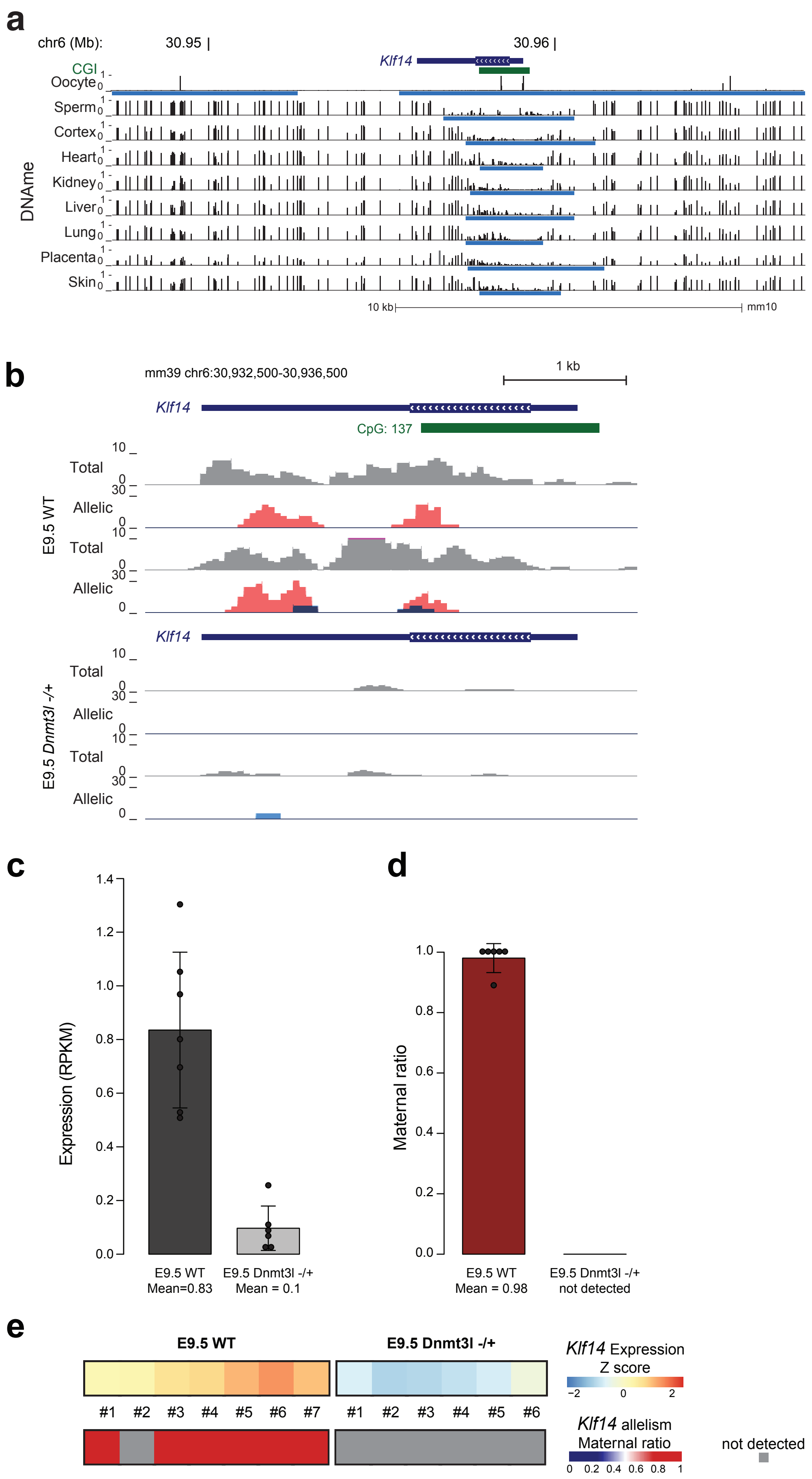

### Supplementary Figure 2

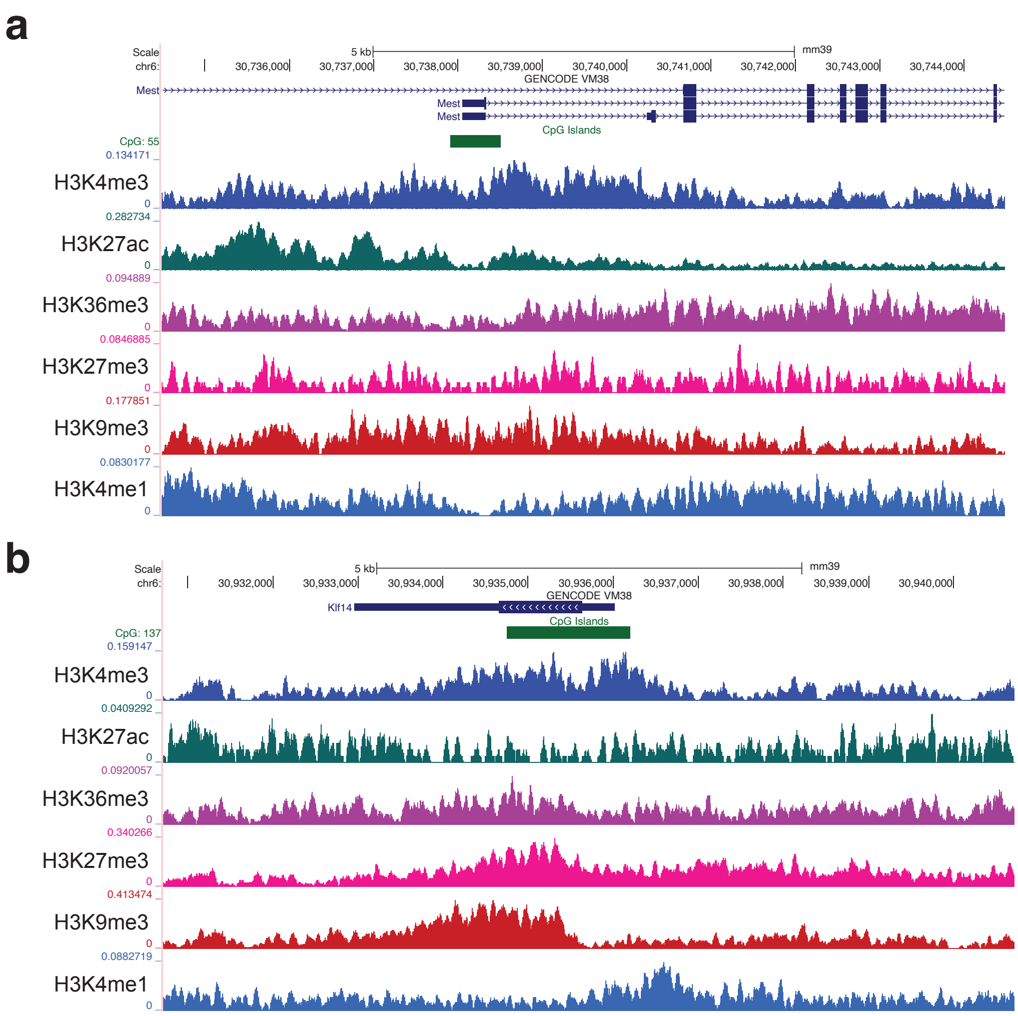

### Supplementary Figure 3

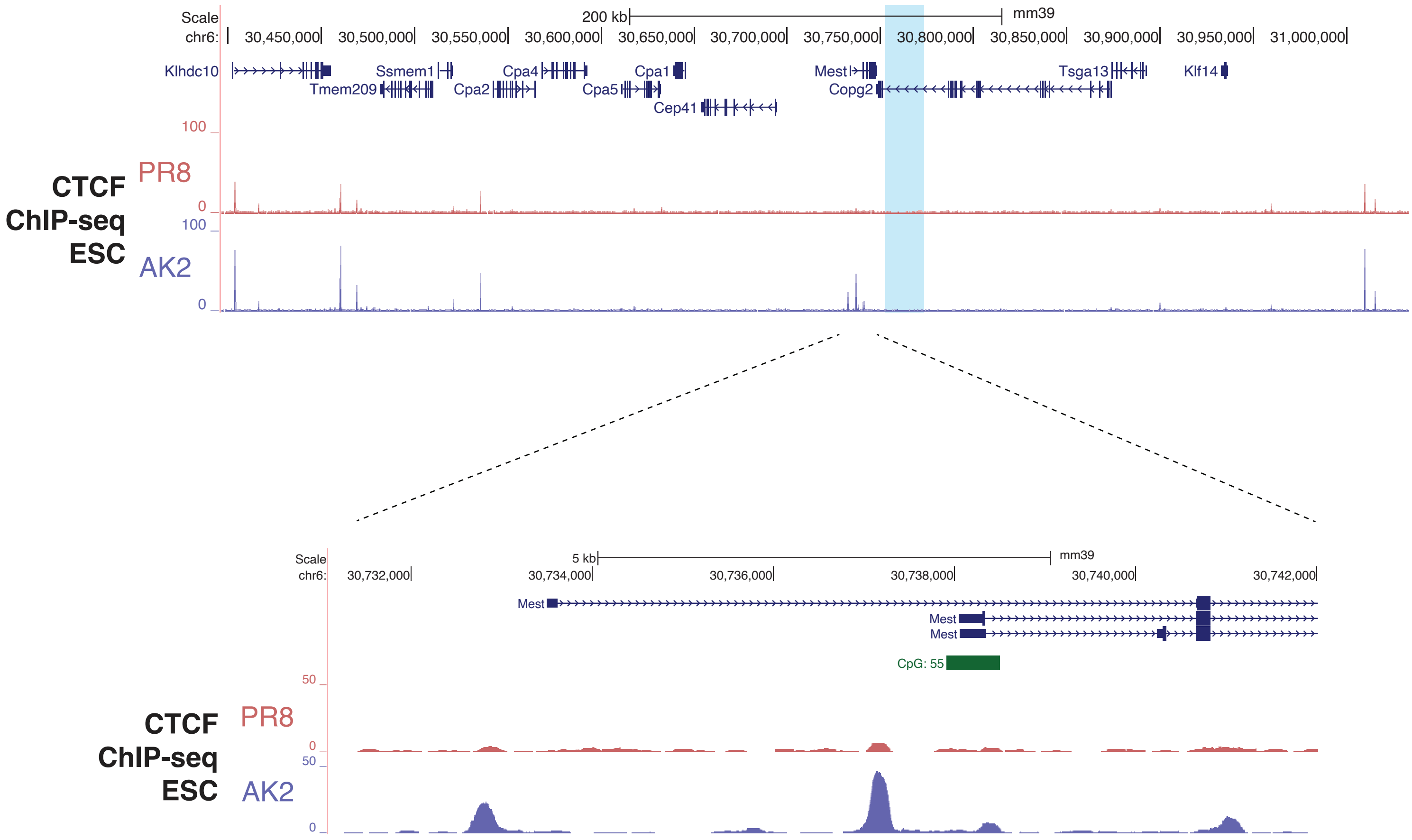

### Supplementary Figure 4

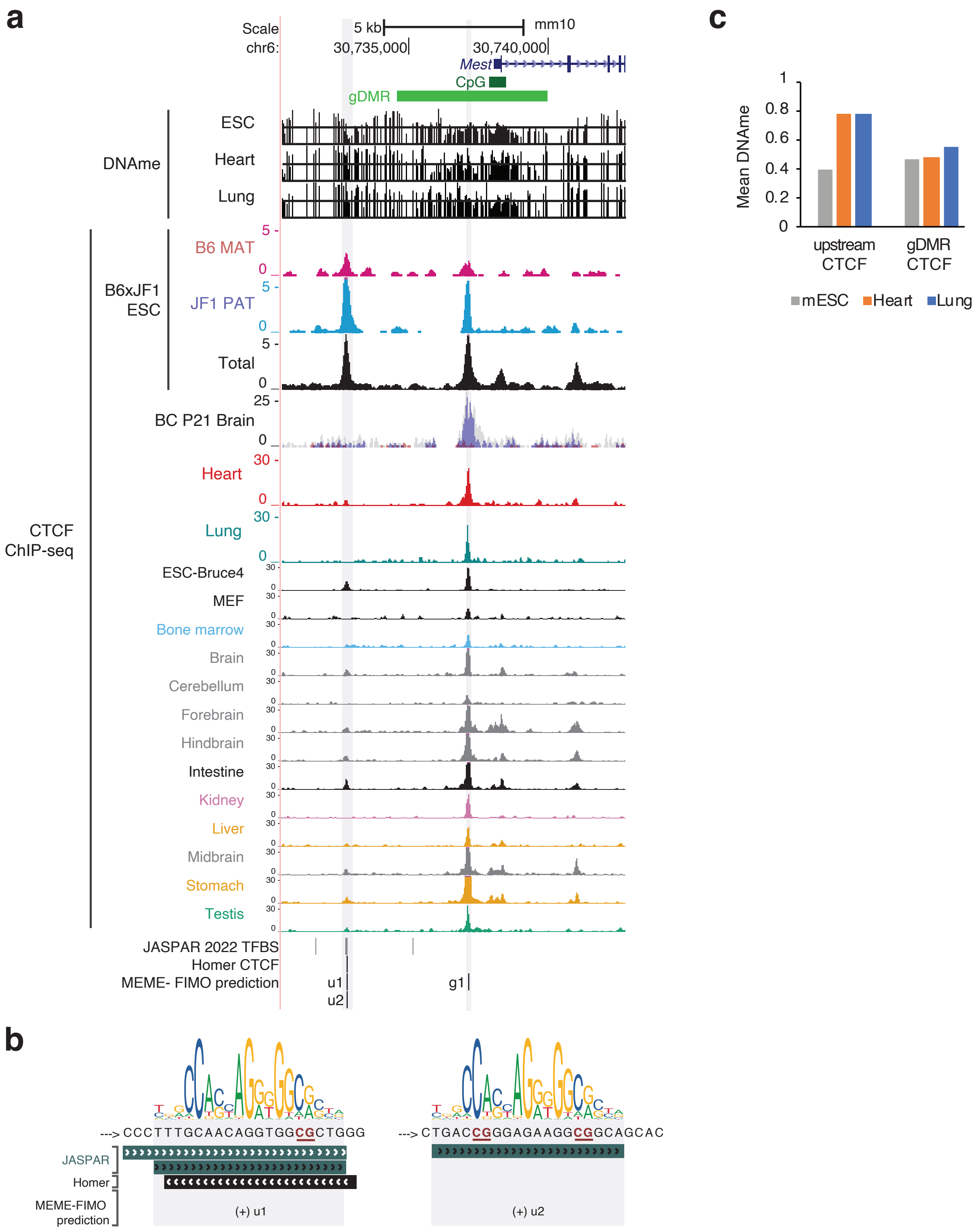

### Supplementary Figure 5

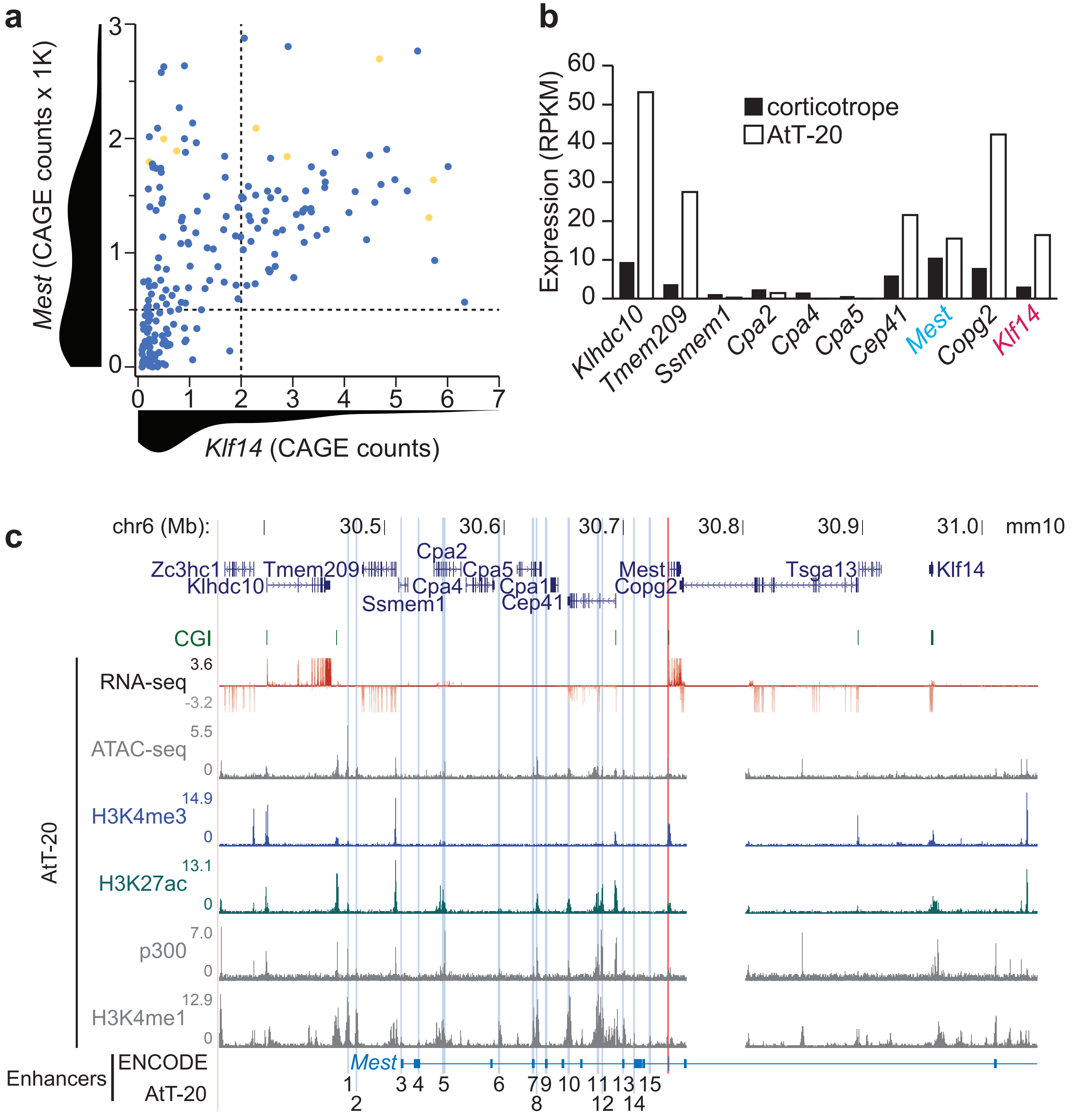

### Supplementary Figure 6

**a**

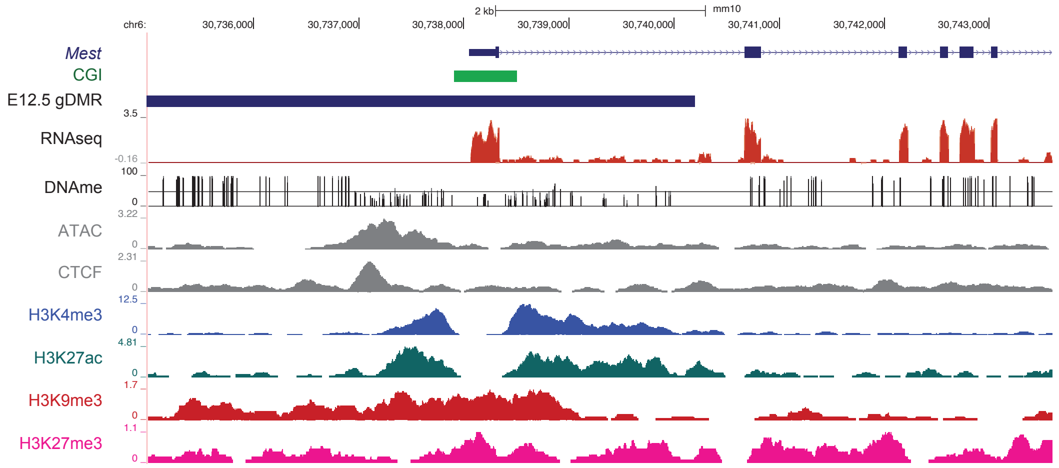


**b**

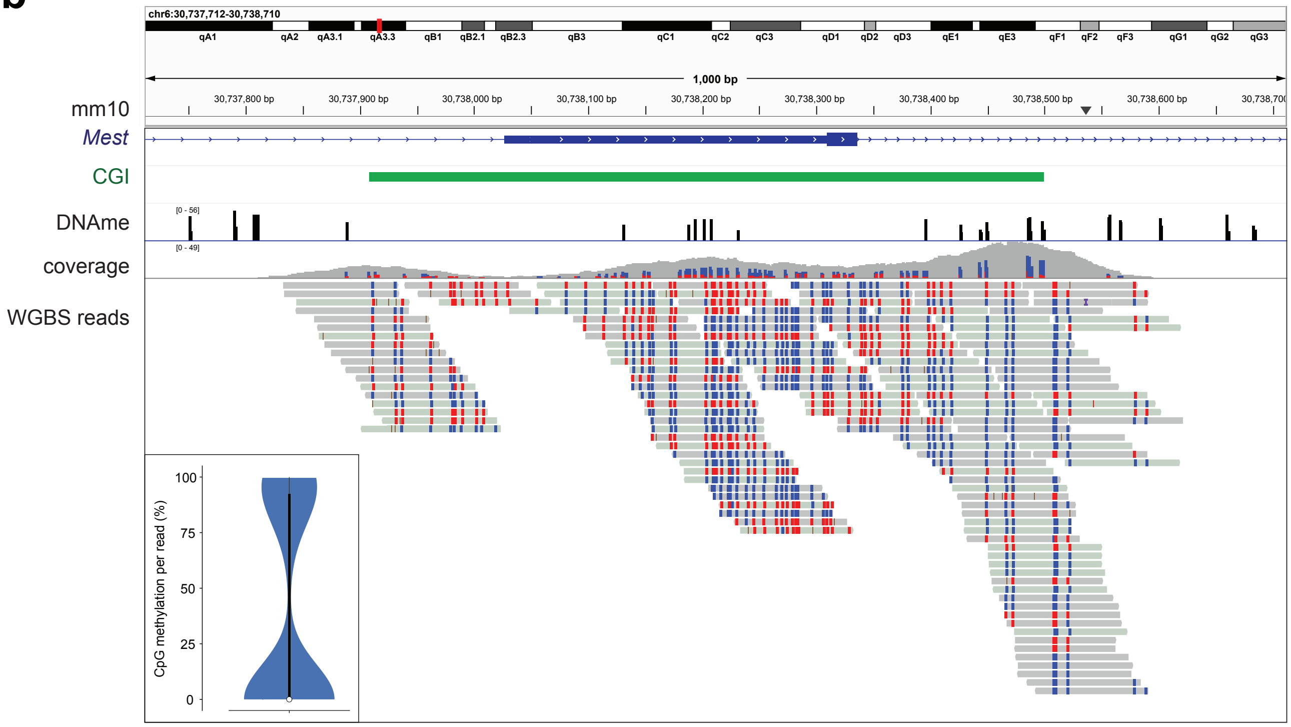


**c**

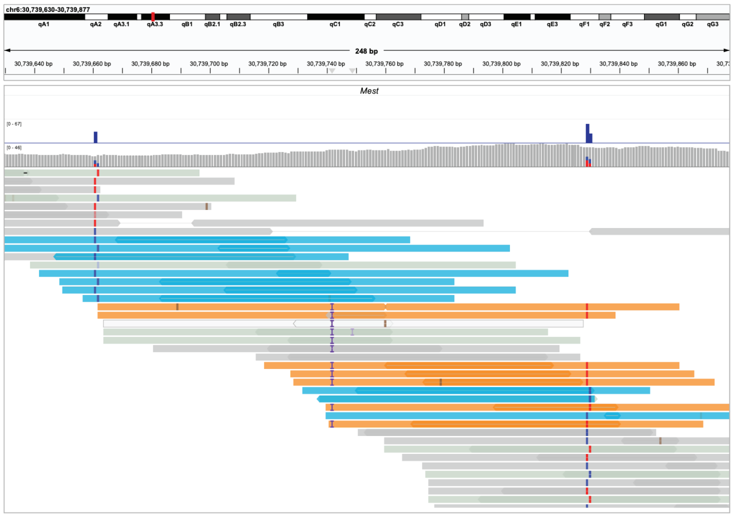

### Supplementary Figure 7

**a**

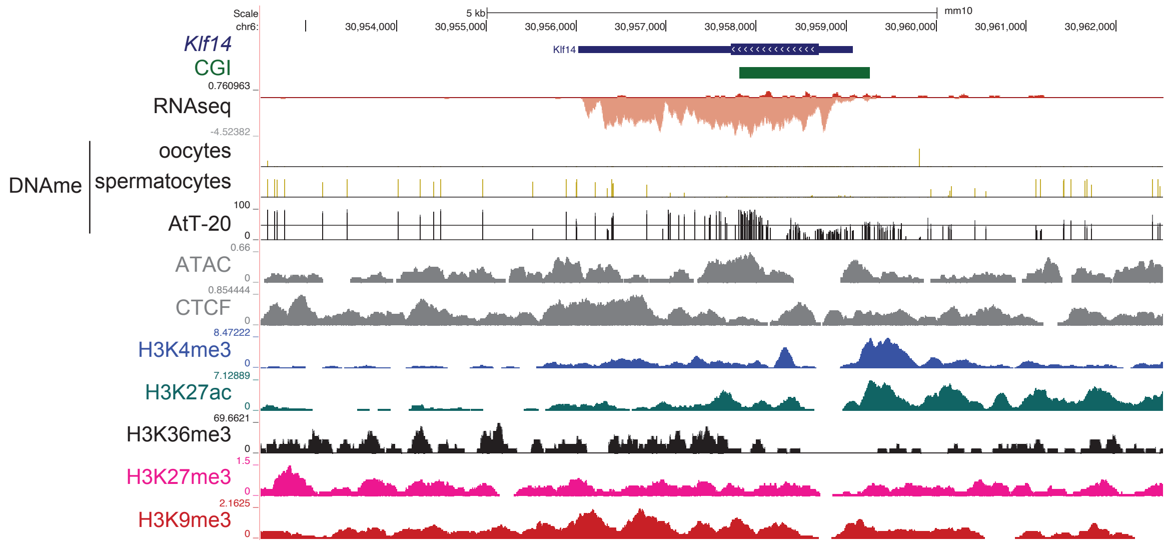


**b**

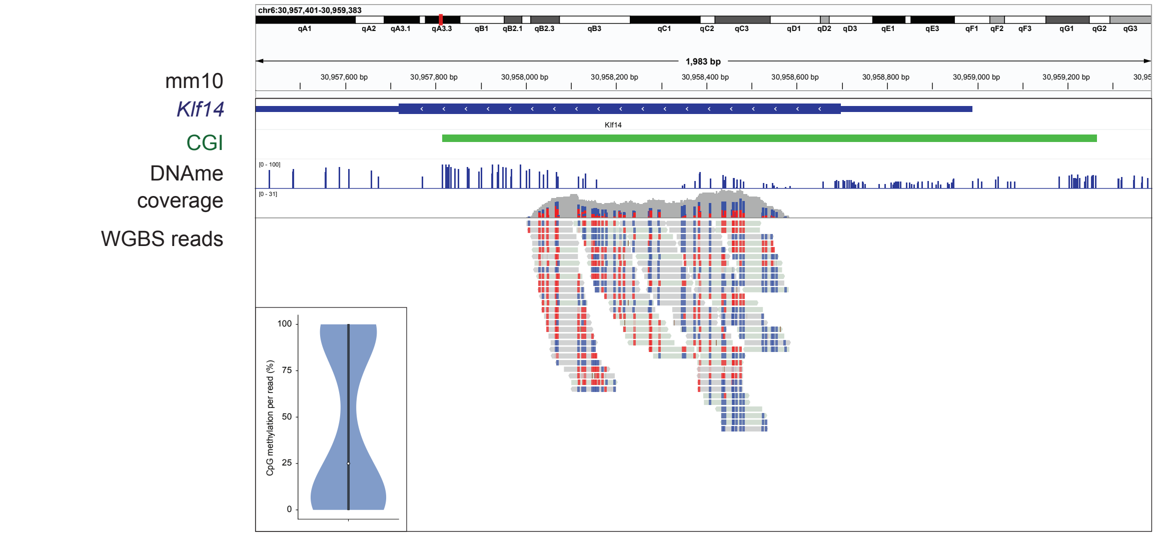

### Supplementary Figure 8

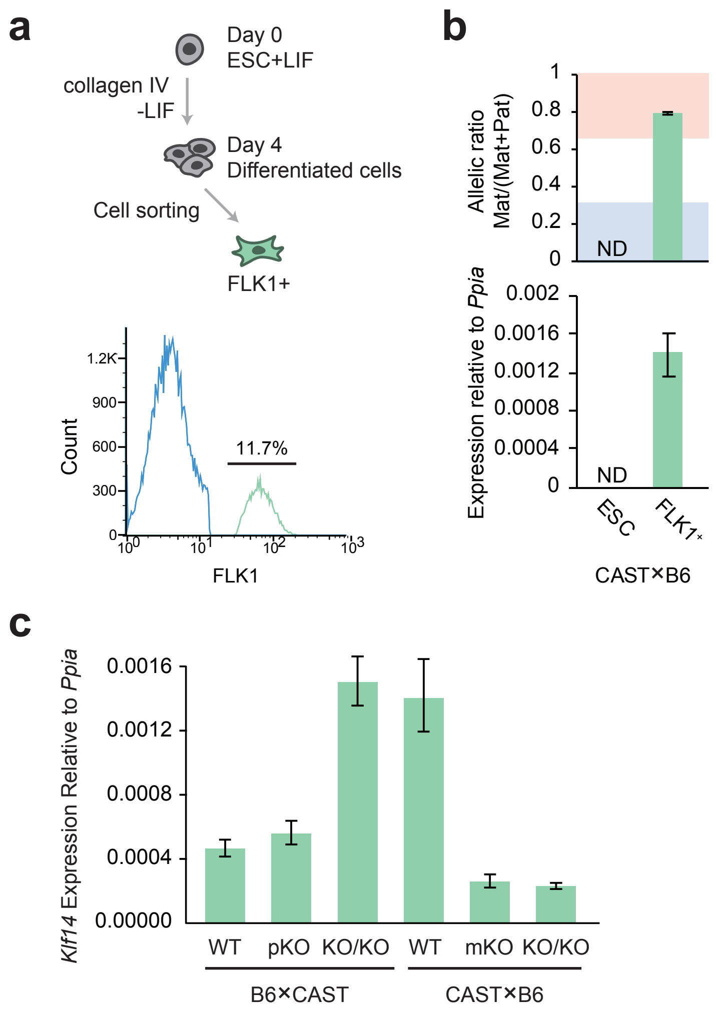

### Supplementary Figure 9

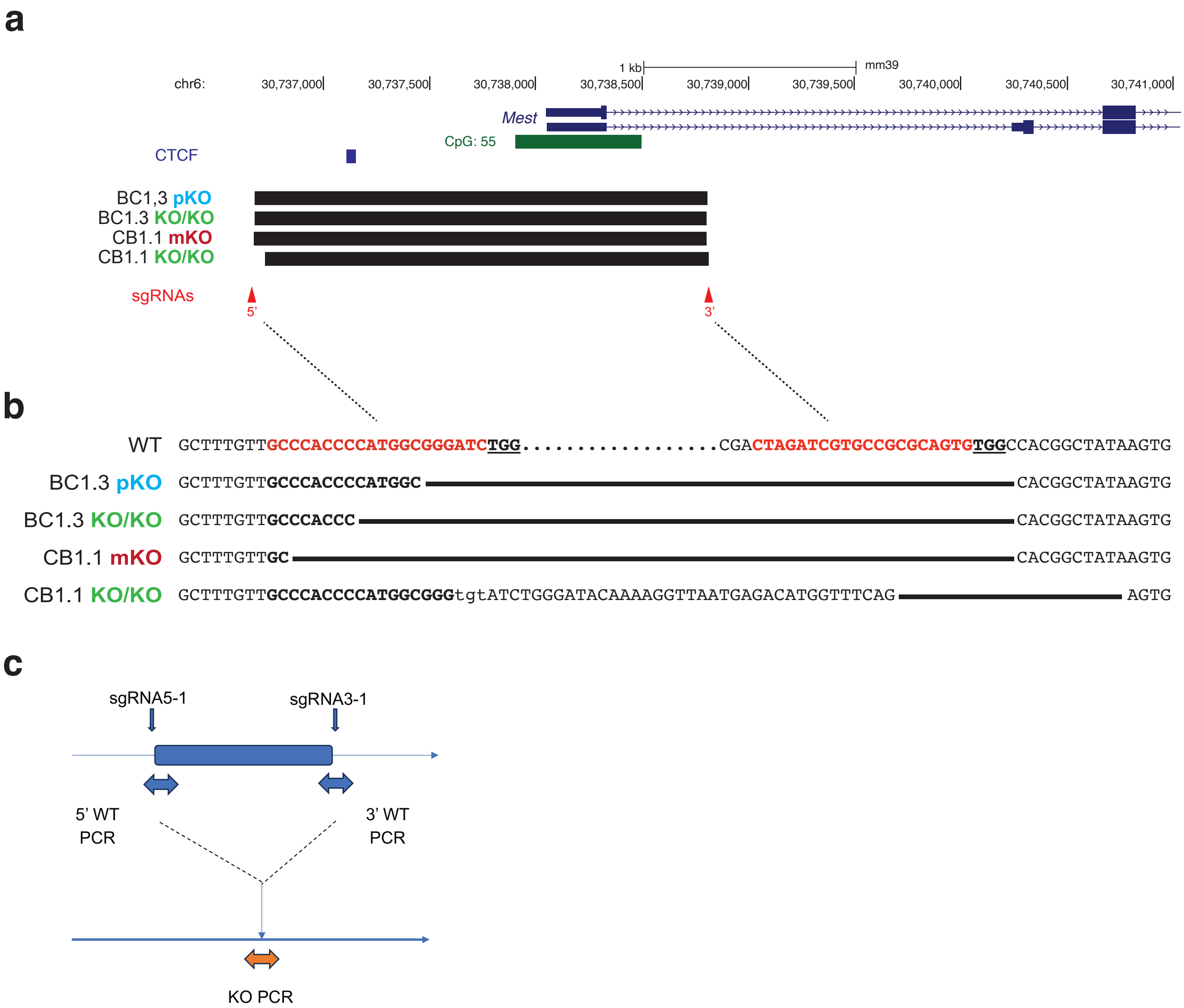


**d**

| Reaction | Forward primer | Reverse primer | Sequencing primer |
|---|---|---|---|
| 5' WT | 5F1 | 5R3 | 5R3 |
| 3' WT | 3F3 | 3R4 | 3F5 |
| proKO | 5F1 | 3R4 | 3F5 |

**e**

| Sequencing primer | BC SNPs monitored |
|---|---|
| 5R3 | rs227043392, rs250442649, rs253971708, rs240077249, rs257640897 |
| 3F5 | rs39519970, rs37511830, rs36857888 |

### Supplementary Figure 10

**a**

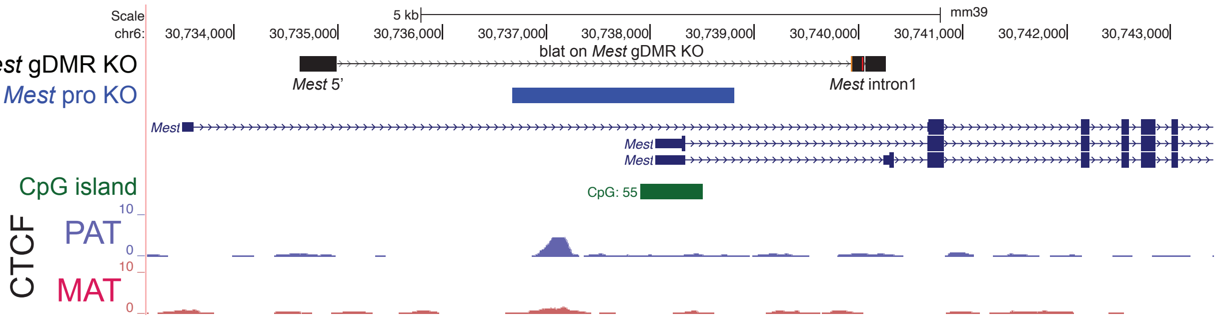


**b**

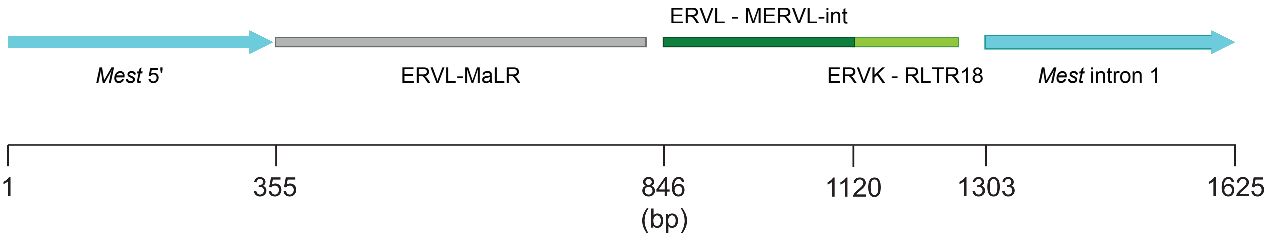

**Supplementary Figure 11.** ***Mest gDMRKO*** **allele in mouse line 48**

~1.6 kb fragment amplified from #48+/del genomic DNA by KN156/157 primers.
Nanopore adaptor attached.
Sequenced using ONT Flongle flowcell.
43 reads were selected, by sequence length and the presence of primer sequences.
A 1625 bp consensus sequence obtained.

>consensus from 43 reads, *Mest* flanking sequences underlined:

```
AGGAGCAAGAGGGTTCAGCTGCTGGCTTCTGCCGCCTTAGCACAGGGCACCCTGAGCTGCTGCTGTT
TCTTGAGGATGAATGGAAAGTCTCACCTAGCCAGTTTCTGTGAAAATGGGGCGTGCTCGGTGTGTGT
GTGTGTGTGTGTGTGTGTGTGTGTGTGTGTATGTGTGTGTGTGTGATATGCCCATAGTAGGAACTTA
AGCTATGCCAGTTGCTTTGCCAGTTGAGTCACATGATTTACAGAAGGTTTTAGTTTTACCCTAAATA
CATTCCTGCATGTCTAAATGTTTCTTAATATCGACTCTGTAGAGCAGAGTACTCTCAGTGCTGCCTG
AGGAGCAAAAAAGCACAGGAAGTTGAAGGAATTGAGCAGAGCAGCTGAGGCTTGGCACTGTGAGAGG
CCATGGAAGGCCATTGGTGAAAGTGCAGCCTCAGTTGCAATTGATGGCCCAGGACTGAAGGGGTCAT
GCAGTGTTTTGGAGATGCCAGTACCATGAGATGACCACCAAGAGCAGCAGCAGCAGTGGAGTAGAGG
CATCTGGAGCCTAGAGGATGACGCGTGTGCTACAAAGGGCATGGCTGGAGAAGTGACCCAAGCCCTT
GGAGGAGCCCAGAAGATCGTGAGTTGGATCCCAGACATTGGACGGTTGGAGATTGATTTTTGCTTTT
GATTGTGACTGTGCCCTGATATTTTCCCTCTTGAAGGAAGAAACTGTTTTAGTGATGCCCACAGTTA
AGAGACTTTTAATTTAAAAAGACTTTGGATTTTTAAAAGAGATATTTTAAAGAGATTGAAATTTTAA
GAATATGTAAAGACTGTGGGACATTTAAAGTTATTTAGACCTCACTCCACTGCTGCTGTGGAGTAGC
ATTGTAACAGGAGTAGAAACCATAGGCATTTGAGCAACTTCTTCATGTAACTTGCTTGTGCCTTCAG
GACCTGCTCTGGCCCGATCACGTATATACCACTTCCATTTGATAATAGACTGCTGCTGTGCGCGTCC
CACTTTATGACTTGCAGGGTCTGATAGTACCCAGCTCATGATGGGTAGTTCAGGTCGCATAGTAACT
TGGTGTCCTATTGTCAAACGTTCAGTTTCCACTAAGGCCCAATAGCAGTTCCAGACAACGGTCAGTC
CAATCAAGACAACGGTCAATCCAATCAAGCTGACTCCAGGGACTCCAGGGACACAGGAAGCATCTAT
ACACAAACTCTTGCAAAAATACAGGAGGCCATGGGGGTCGATTCTAGAGCTCCACTGCTAAACTCCA
CTGCTGCTGCTGCTCTTGGTGGTCACCTCAAAGGAAATGGTTTCTAATCGTGCCTTATTTCTCGCCT
CCAGGATAAATTGATCTGGACCTTTGAAATCCAGGCATGGCAAGCTGGGGCCACAAATCTGTGTTCC
CCCCCTCCTGCCCCCCCCCCACAAAAGTCTTGAGTGCTAGGTGTGGGGGTGGGGTCTTAATTAAATA
AAAGGGAACCACCTCAAGAGTCTGCTAAGAATGCACAAATATAATCTGCTGCCCAGACTTGTAAGAG
ATTAGATATTCAGGGTCAGGGCAAAGCTGCTTACTGGAATTCTCCTGGTCTCTCCCCTCAACAGTTC
CATAGCCCAAATGCAGA
```

>consensus from 43 reads masked and annotated

AGGAGCAAGAGGGTTCAGCTGCTGGCTTCTGCCGCCTTAGCACAGGGCACCCTGAGCTGCTGCTGTT
TCTTGAGGATGAATGGAAAGTCTCACCTAGCCAGTTTCTGTGAAAATGGGGCGTGCTCGGTGTGTGT
GTGTGTGTGTGTGTGTGTGTGTGTGTGTGTATGTGTGTGTGTGTGATATGCCCATAGTAGGAACTTA
AGCTATGCCAGTTGCTTTGCCAGTTGAGTCACATGATTTACAGAAGGTTTTAGTTTTACCCTAAATA
CATTCCTGCATGTCTAAATGTTTCTTAATATCGACTCTGTAGAGCAGAGTACTCTCAGTGCTGCCTG
AGGAGCAAAAAAGCACAGGAAGTNNNNNNNNNNNNNNNNNNNNNNNNNNNNNNNNNNNNNNNNNNNN
NNNNNNNNNNNNNNNNNNNNNNNNNNNNNNNNNNNNNNNNNNNNNNNNNNNNNNNNNNNNNNNNNNN
NNNNNNNNNNNNNNNNNNNNNNNNNNNNNNNNNNNNNNNNNNNNNNNNNNNNNNNNNNNNNNNNNNN
NNNNNNNNNNNNNNNNNNNNNNNNNNNNNNNNNNNNNNNNNNNNNNNNNNNNNNNNNNNNNNNNNNN
NNNNNNNNNNNNNNNNNNNNNNNNNNNNNNNNNNNNNNNNNNNNNNNNNNNNNNNNNNNNNNNNNNN
NNNNNNNNNNNNNNNNNNNNNNNNNNNNNNNNNNNNNNNNNNNNNNNNNNNNNNNNNNNNNNNNNNN
NNNNNNNNNNNNNNNNNNNNNNNNNNNNNNNNNNNNNNNNNNNNNNNNNNNNNNNNNNNNNNNNNNN
NNNNNNNNNNNNNNNNNNNNNNNNNNNNNNNNNNNNNNNNNNCACTCCACTGCTGCTGTGGAGTANN
NNNNNNNNNNNNNNNNNNNNNNNNNNNNNNNNNNNNNNNNNNNNNNNNNNNNNNNNNNNNNNNNNNN
NNNNNNNNNNNNNNNNNNNNNNNNNNNNNNNNNNNNNNNNNNNNNNNNNNNNNNNNNNNNNNNNNNN
NNNNNNNNNNNNNNNNNNNNNNNNNNNNNNNNNNNNNNNNNNNNNNNNNNNNNNNNNNNNNNNNNNN
NNNNNNNNNNNNNNNNNNNNNNNNNNNNNNNNNNNNNNNNNNNNNNNNNTTCCAGACAACGGTCAGT
CCAATCAAGACAACGGTCAATCCAATCAAGCTGACTCCAGGGACTCCAGGGACACAGGAAGCATCTA
TACACAAACTCTTGCAAAAATACAGGAGGCCATGGGGGTCGATTCTAGAGCTCCACTGCTAAACTCC
ACTGCTGCTGCTGCTCTTGGTGGTCACCTCAAAGGAAATGGTTTCTAATCGTGCCTTATTTCTCGCC
TCCAGGATAAATTGATCTGGACCTTTGAAATCCAGGCATGGCAAGCTGGGGCCACAAATCTGTGTTC
CCCCCCTCCTGCCCCCCCCCCACAAAAGTCTTGAGTGCTAGGTGTGGGGGTGGGGTCTTAATTAAAT
AAAAGGGAACCACCTCAAGAGTCTGCTAAGAATGCACAAATATAATCTGCTGCCCAGACTTGTAAGA
GATTAGATATTCAGGGTCAGGGCAAAGCTGCTTACTGGAATTCTCCTGGTCTCTCCCCTCAACAGTT
CCATAGCCCAAATGCAG

Green: Primer sequences (KN156/157)
Lightblue: Mest-DMR flanking regions
NNNNNNNNN: Family: ERVL-MaLR
NNNNNNNNN: Family: ERVL (MERVL-int)
Chr13: Chr13 sequence (older ERVK element, RLTR18)

### Supplementary Figure 12

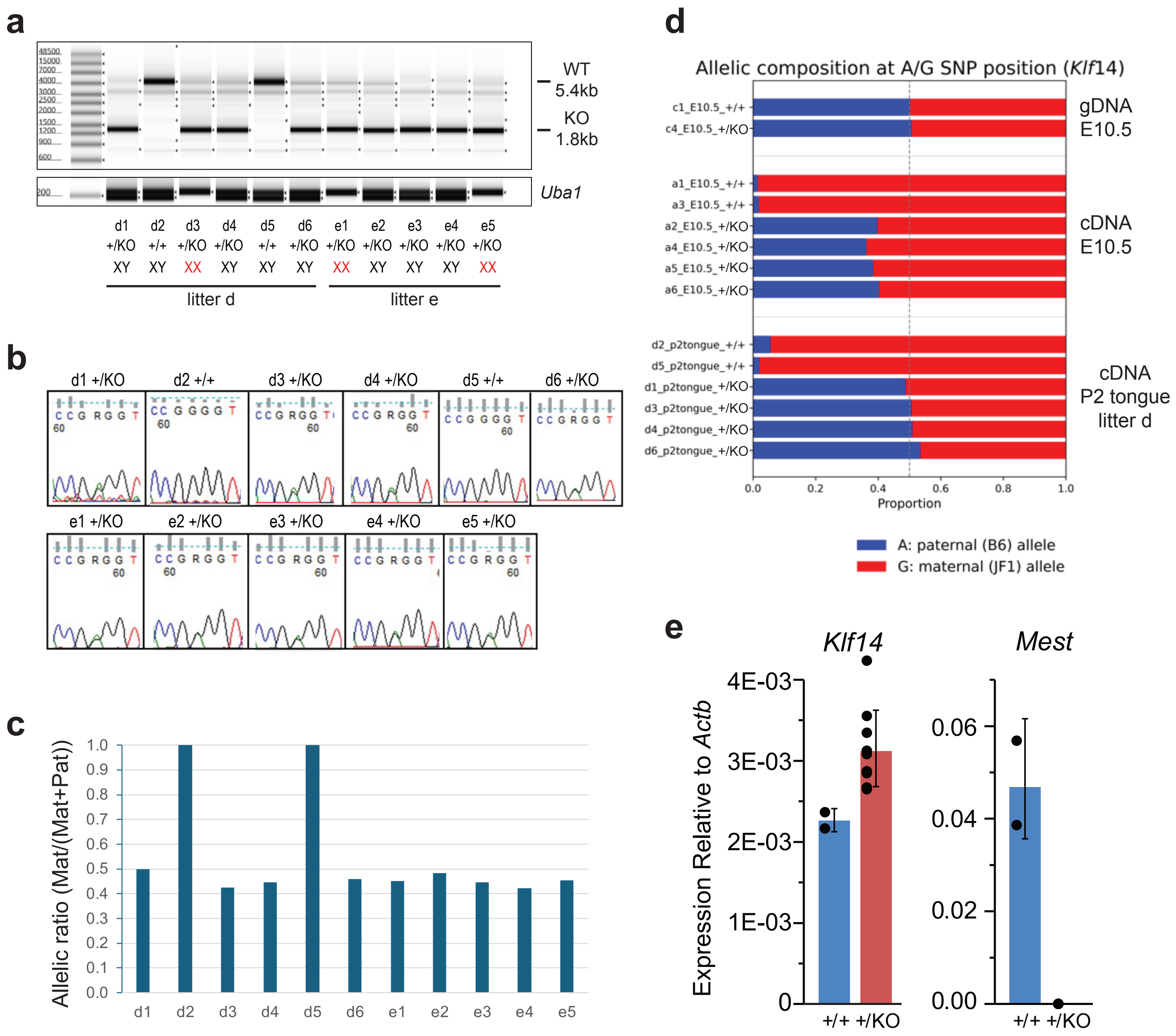

### Supplementary Figure 13

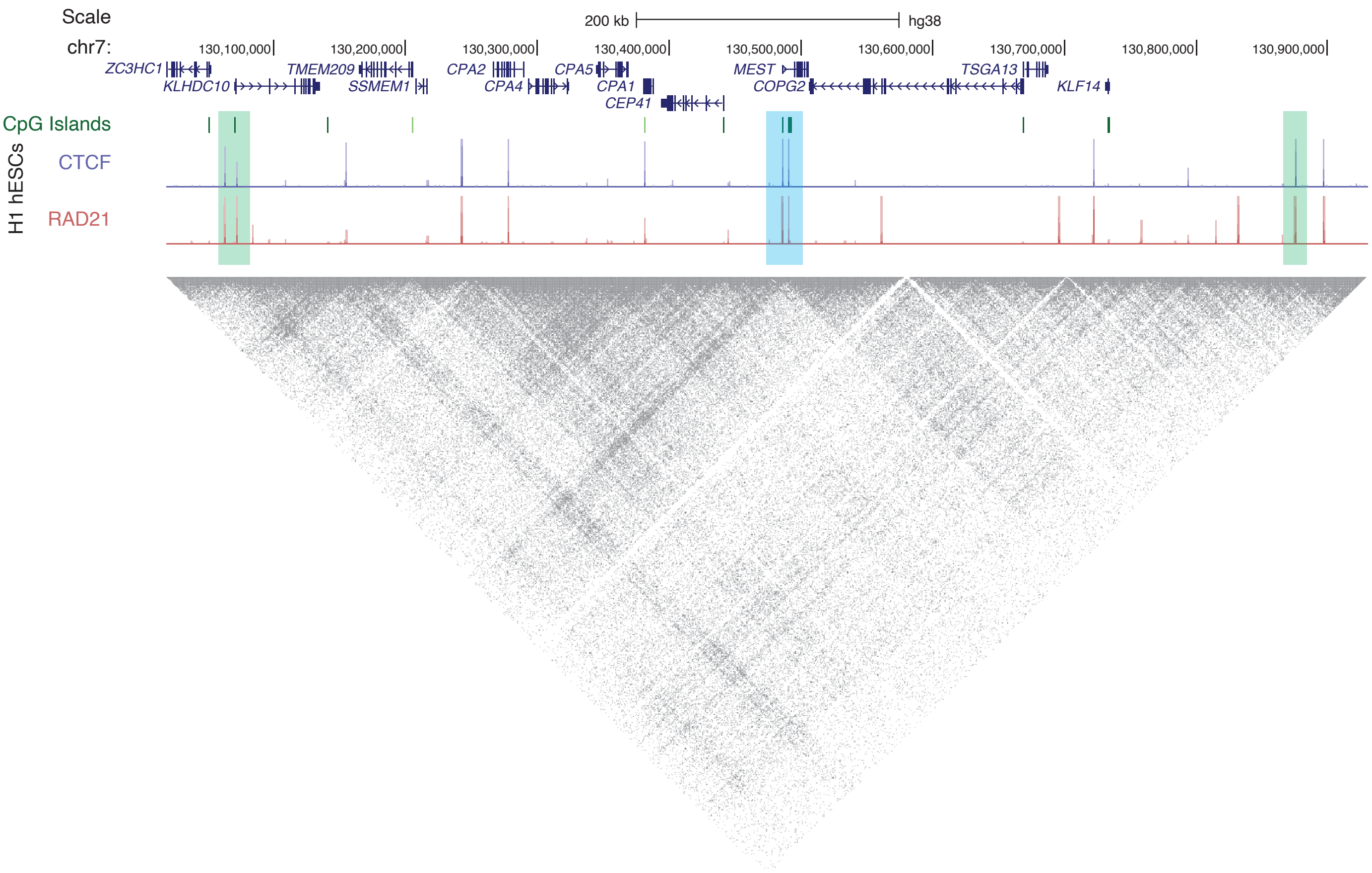

**Supplementary Table 1. Datasets used in this study**

| Species | Tissue | F1 crosses | Data type | Accession | Reference |
|---|---|---|---|---|---|
| Mouse | J1 ESC | | DNAme | GSE61457 | Li, Z. *et al. Genome Biol* 16:115 (2015) |
| Mouse | ESC | B6×CAST | ZFP57 ChIP-seq | GSE55382. | Strogantsev, R. *et al. Genome Biol* 16:112 (2015) |
| Mouse | Brain | B6×CAST | CTCF ChIP-seq | GSE35140 | Prickett, A. *et al. Genome Res* 23:1624–1635 (2013) |
| Mouse | ICM | B6X×PWK | H3K4me3 | GSE71434 | Zhang, B. *et al. Nature* 537:553–557 (2016) |
| Mouse | ICM | B6X×PWK | H3K27me3 | GSE76687 | Zheng, H. *et al. Mol Cell* 63(6):1066-1079 (2016) |
| Mouse | ICM | B6×DBA | H3K9me3 | GSE98149 | Wang, C. *et al. Nat Cell Biol* 20(5):620-631 (2018) |
| Mouse | Various F1 tissues | CAST×FVB | RNA-seq | GSE75957 | Andergassen, D. *et al. eLife* 6:e25125 (2017) |
| Mouse | Adult heart, lung | | DNAme | GSE42836 | Hon G.C., *et al. Nat Genet* 45(10):1198-206 (2013) |
| Mouse | Adult heart, lung | | CTCF ChIP-seq | GSE49847 | Yue, F. *et al. Nature* 515:355-64 (2014) |
| Mouse | BJ1, JB1 | B6X×JF1 | CTCF ChIP-seq | GSE207166 | Farhadova, S. *et al. Nucleic Acids Res.* 52, 6183–6200 (2024) |
| Mouse | AK2, PR8 mESC | | CTCF ChIP-seq | PRJEB28762 | Llères, D. *et al. Genome Biol* 20:272 (2019) |
| Mouse | Various | | CTCF ChIP-seq | ENCODE | Absacal, F. *et al. Nature* 583:699-710 (2020) |
| Mouse | Various | | DNAme | DRA000484 | Kobayashi, H. *et al. PLoS Genet* 8(1):e1002440 (2012) |
| Mouse | Various | | DNAme | MethBase | Song, Q. *et al. PLoS ONE* 8(12): e81148 (2013) |
| Mouse | differentiated ESC | | RNA-seq | GSE69101 | Goode, D.K. *et al. Dev Cell* 36:572-587 (2016) |
| Mouse | ESC | | Hi-C | GSM2533821<br>GSM2533820<br>GSM2533819<br>GSM2533818 | Bonev, B. *et al. Cell* 171:557-572 (2017) |
| Mouse | NPC | | ChIP-seq | GSE96107 | Bonev, B. *et al. Cell* 171:557-572 (2017) |
| Mouse | AtT-20 | C57L/J×A/J | WGBS, ChIP-seq | GSE87185 | Mayran, A. *et al. Nat Genet* 50(2):259-269 (2018) |
| Human | H1 hESCs | | Micro-C | | Krietenstein, N. *et al. Molecular Cell* 78:554-565 (2020) |

**Supplementary Table 2. Putative *Mest* enhancers in AtT-20 cells**

| | | mm10 | | mm39 | |
|---|---|---|---|---|---|
| **Enh name** | **chr** | **start** | **end** | **start** | **end** |
| AtT20-E1 | chr6 | 30469398 | 30469763 | 30469397 | 30469762 |
| AtT20-E2 | chr6 | 30476424 | 30477698 | 30476423 | 30477697 |
| AtT20-E3 | chr6 | 30514205 | 30514570 | 30514204 | 30514569 |
| AtT20-E4 | chr6 | 30528835 | 30529047 | 30528834 | 30529046 |
| AtT20-E5 | chr6 | 30548932 | 30551641 | 30548931 | 30551640 |
| AtT20-E6 | chr6 | 30595521 | 30597461 | 30595520 | 30597460 |
| AtT20-E7 | chr6 | 30623888 | 30625866 | 30623887 | 30625865 |
| AtT20-E8 | chr6 | 30627634 | 30628614 | 30627633 | 30628613 |
| AtT20-E9 | chr6 | 30635198 | 30636735 | 30635197 | 30636734 |
| AtT20-E10 | chr6 | 30653782 | 30655803 | 30653781 | 30655802 |
| AtT20-E11 | chr6 | 30678280 | 30679931 | 30678279 | 30679930 |
| AtT20-E12 | chr6 | 30682239 | 30683560 | 30682238 | 30683559 |
| AtT20-E13 | chr6 | 30699917 | 30701069 | 30699916 | 30701068 |
| AtT20-E14 | chr6 | 30708865 | 30709899 | 30708864 | 30709898 |
| AtT20-E15 | chr6 | 30721754 | 30723284 | 30721753 | 30723283 |

**Supplementary Table 3. 3C-qPCR primers and results**

| Primer Name | Primer sequence (5'-3') | Distance from *Mest* promoter (bp) | Relative contact frequency | |
|---|---|---|---|---|
| | | | Values (A.U.) | S.E.M. |
| **1** | ggaaagcaaaaggcattgttcaactcc | -318866 | 0.7250865 | 0.1980686 |
| **3** | gaaagagtgaccaggaaagcctc | -309339 | 0.9299781 | 0.2488751 |
| **7F** | gggggtgtcatgcttttctttgag | -297421 | 1.1497746 | 0.2865521 |
| **7R** | gctgttttcccctcgcctatac | -288439 | 0.8769321 | 0.5104782 |
| **8** | cccccaaagtgaaaggctctatg | -285720 | 0.9198293 | 0.2453514 |
| **9** | gctctatctcccagaccaagag | -280436 | 0.5250920 | 0.4166086 |
| **11** | ggacatcagactgcagaaggtg | -277394 | 0.4630266 | 0.1663155 |
| **12** | gtagctctgcaacagagtcctg | -274605 | 1.0755548 | 0.3665478 |
| **13** | cctcgccctaactttccaagtc | -271217 | 1.3762919 | 0.2795983 |
| **16** | tttcccctccttgtcatggctc | -261325 | 1.3633331 | 0.4766273 |
| **17** | caaccactaagctacagcacctg | -255575 | 0.9461413 | 0.4036599 |
| **19** | gtggtgtgcttgggagatagag | -243500 | 0.8741589 | 0.3102070 |
| **20** | gagaagtcacctgacaggtcac | -242839 | 0.8906782 | 0.3165653 |
| **24** | gcatgcctccaaattgtggctg | -229821 | 0.9818680 | 0.2296669 |
| **26** | cccgtctctctcttgactcttc | -222259 | 1.9326535 | 1.0311161 |
| **27** | tatgttgagggccacagccttg | -217955 | 2.0854678 | 0.8843397 |
| **28** | gtcaggaggtaatgtcccacac | -215366 | 1.9001838 | 0.9062894 |
| **31F** | cagtgctttatgagcagcaggtg | -205672 | 0.9720767 | 0.0522702 |
| **31R** | aagaggtgtacctgctcctagc | -187642 | 2.2037396 | 1.1676301 |
| **32** | ccggagaattcagagttccctg | -183321 | 1.2715486 | 0.8905498 |
| **33** | gagaaacaggtttcaggtagtacgtc | -182670 | 1.2610068 | 0.7614388 |
| **36** | ggctggatcagtcccataatcag | -180798 | 1.2893922 | 0.1543019 |
| **43** | ggtcctcagcttctgtgcttag | -171501 | 1.3680677 | 0.3741725 |
| **45** | ggagatggttcagtgggcaaac | -141040 | 1.3322483 | 0.4028421 |
| **47F** | caggacctgtggaactcaagag | -138895 | 2.3220714 | 0.9125384 |
| **47R** | gcccagccttaggctagttttaac | -135163 | 2.9262374 | 1.0514041 |
| **49** | tgatgctgtaggcaaggccatg | -116380 | 1.3490650 | 0.4140272 |
| **50** | tagtactagagctgagcccgtg | -114190 | 0.7142543 | 0.1259559 |
| **52** | ccagaagcaacaggcttgagag | -108996 | 1.4856598 | 0.8292313 |
| **53** | gcaggagtagaaaacagcccag | -105382 | 1.1912683 | 0.1757772 |
| **54** | actgggcgtgtgagcagattag | -104939 | 1.5106957 | 0.3013021 |
| **55** | cctgccccacttgatgaatgac | -103198 | 1.8492005 | 0.6555346 |
| **58** | gaaccaaaggagctacctgcag | -101002 | 1.4042957 | 0.5444066 |
| **59** | cagaggaatcctgagtgggaag | -100433 | 0.9337705 | 0.3834238 |
| **61** | gctggctggctggtttgttttc | -94109 | 1.1229046 | 0.3802015 |
| **62** | atgacagatggtggggacagag | -89445 | 1.6250945 | 0.6210862 |
| **65** | aatcgggactgaagtggaggtc | -81500 | 2.4841154 | 1.5209152 |
| **66** | ctgtgactgtggataagagggac | -81089 | 1.5088341 | 0.3532613 |
| **70** | ggggtcctgtctccaacaacag | -68943 | 1.5802145 | 1.0222665 |
| **75** | cctgtagttagtcggagacagag | -53916 | 7.0268250 | 1.4071487 |
| **78F** | cacaggtggtgagatagcctac | -41082 | 2.8890247 | 1.0645041 |
| **84** | aaaagctcctgggagcccatac | -31292 | 1.1350646 | 0.4657182 |
| **86** | gctaccttgtcttgctccaagc | -25177 | 4.3312487 | 1.5470322 |
| **88F** | gatgtctcctgttctgctggag | -22947 | 10.6703605 | 4.4154437 |
| **88R** | gctgcttgagtcggcaaataatgg | -22153 | 7.4254610 | 2.2211650 |
| **89** | cggaatttgggagatgtagctaagg | -18068 | 1.9633990 | 1.0855409 |
| **91** | ctaaaaacgtggcttggggagtg | -12198 | 3.6838559 | 1.4274761 |
| **92** | catggttctgatggccaggctc | -7078 | 3.4660960 | 1.1067692 |
| **94** | gtgccagaccctgtctatatcc | 0 | **Bait primer (*Mest* promoter)** | |

**Supplementary Table 4.1 Chromosome count and sex of new F1 ESC lines**

| Female | Male | Line | Sex | % 40 chromosomes |
|---|---|---|---|---|
| C57BL/6 | CAST | BC1.1 | XY | 93 |
| | | BC1.2 | XX | ND |
| | | BC1.3* | XY | 95 |
| | | BC2.1 | XY | 88 |
| | | BC2.2 | XX | ND |
| | | BC2.3 | XY | 91 |
| | | BC2.4 | XX | ND |
| CAST | C57BL/6 | CB1.1* | XY | 90 |
| | | CB1.2 | XX | ND |
| | | CB1.3 | XX | 95 |

*F1 ESC lines used in this study
ND: not determined

**Supplementary Table 4.2 *Mest* imprinted expression in F1 ESCs**

| | | Allelic ratio Mat/(Mat+Pat) |
|---|---|---|
| **BC 1.3** | gDNA | 0.476 |
| | cDNA | 0.239 |
| **CB 1.1** | gDNA | 0.530 |
| | cDNA | 0.222 |

**Supplementary Table 5. Primer sequences used in this study**

| Primer name | Sequence (5'-3') | Reaction | Reference |
|---|---|---|---|
| **RT-qPCR** | | | |
| Ppia F1 | CGCGTCTCCTTCGAGCTGTTTG | *Ppia* | Mamo et al., 2007 |
| Ppia R1 | TGTAAAGTCACCACCCTGGCACAT | *Ppia* | Mamo et al., 2007 |
| Klf14 qF1 | GGTGAGAGCAGCTAGACAATAG | *Klf14* | This study |
| Klf14 qR1 | CTGGGACACACAACAGAAGA | *Klf14* | This study |
| KN250 | GCTGTGTCCCAAGCAGTTCT | *Klf14* | This study |
| KN251 | GGTTGTGCTTGAGGGAAAGA | *Klf14* | This study |
| KN341 | GCTTCACTGCTTGCCTGTAG | *Klf14* | This study |
| KN342 | TTTGTCTGAACCCATCCCCA | *Klf14* | This study |
| KN252 | GATTCGCAACAATGACGGCA | *Mest* | This study |
| KN253 | ATCCAGAATCGACACTGTGG | *Mest* | This study |
| Actb-F | CCAACTGGGACGACATGG | *Actb* | This study |
| Actb-R | GGGCACAGTGTGGGTGAC | *Actb* | This study |
| **Pyrosequencing** | | | |
| Biotin-Klf14 F3 | GATGAGCTAGCCCGCCACT | *Klf14* | This study |
| Klf14 R1 | GGCCAGAAGAGGAGCTGGAG | *Klf14* | This study |
| Klf14 S1 | AGTGCGACGACGACC | *Klf14* | This study |
| Klf14 S2 | TTTTCACCGGTGTGCGT | *Klf14* | This study |
| ***Mest proKO* sgRNA cloning** | | | |
| Sap1-sgRNA-5-1 | TGTGGAAAGGGCTCTTCACCGcccaccccatggcgggatcGTTTTAGAGCTAGAAATAGCAAGTTAAAATAAGGCTAG | sgRNA-5-1 F | This study |
| Sap1-sgRNA-3-1 | TGTGGAAAGGGCTCTTCACCGctagatcgtgccgcgcagtgGTTTTAGAGCTAGAAATAGCAAGTTAAAATAAGGCTAG | sgRNA-3-1 F | This study |
| XbaI-sgRNA | ggcgcacgcactaAAAAAtcgTCTAGAAAAAAAGCACCGACTCGGTGCCACTTTTTCAAGTTGATAACGGACTAGCCTTATTTTAACTTGCTATTTCTAG | sgRNA R | This study |
| ***Mest proKO* genotyping** | | | |
| Mest-5F1 | CCTTCCTCCCTCCCCTTAAT | *Mest* | This study |
| Mest-5R3 | CAGGCTGCGATCACATAGAA | *Mest* | This study |
| Mest-3F3 | TCACAGAGCGACTTAAAATTTCC | *Mest* | This study |
| Mest-3R4 | AGTGCCAGAACACGATGGTT | *Mest* | This study |
| Mest-3F5 | GTGTGGCACCTCTGTTTCC | *Mest* | This study |
| ***Mest gDMRKO* sgRNA cloning** | | | |
| sgRNA1-F | AGCAAAAAAGCACAGGATTGTTTTAGAGCTAGAAATAGCAAG | sgRNA1 | This study |
| sgRNA1-R | AACAATCCTGTGCTTTTTTGCTCGGTGTTTCGTCCTTTCCAC | sgRNA1 | This study |
| sgRNA2-F | TGACCATGGCGGGTCACAAGTTTTAGAGCTAGAAATAGCAAG | sgRNA2 | This study |
| sgRNA2-R | AACTTGTGACCCGCCATGGTCACGGTGTTTCGTCCTTTCCAC | sgRNA2 | This study |
| **sgRNA in vitro templates** | | | |
| T7-sgRNA1-F | ttaatacgactcactataggAGCAAAAAAGCACAGGATT | sgRNA1 | This study |
| T7- sgRNA2-F | ttaatacgactcactataggTGACCATGGCGGGTCACAA | sgRNA2 | This study |
| gRNA-R | AAAAGCACCGACTCGGTGCC | sgRNA | Inui *et al.* 2014 |
| ***Mest gDMRKO* genotyping** | | | |
| KN156 | AGGAGCAAGAGGGTTCAGC | *Mest* | This study |
| KN157 | CTGCATTTGGGCTATGGAAC | *Mest* | This study |
| **hCas9 template PCR** | | | |
| T7 Cas9 Fw | taatacgactcactatagggAGAATGGACAAGAAGTACTCCATTGG | *hCAs9* | Inui *et al.* 2014 |
| Cas9 Rv | TCACACCTTCCTCTTCTTC | *hCas9* | Inui *et al.* 2014 |
| **4C-distal viewpoint** | | | |
| Klf14-19 F1-P7 2.1 | CAAGCAGAAGACGGCATACGAGATCGTGATGTGACTGGAGTTCAGACGTGTGCTCTTCCGATCTttttcagctcctcctgaacc | 4C-seq *Klf14* distal | This study |
| Klf14-19 F1-P7 2.2 | CAAGCAGAAGACGGCATACGAGATACATCGGTGACTGGAGTTCAGACGTGTGCTCTTCCGATCTttttcagctcctcctgaacc | 4C-seq *Klf14* distal | This study |
| Klf14-19 F1-P7 2.4 | CAAGCAGAAGACGGCATACGAGATTGGTCAGTGACTGGAGTTCAGACGTGTGCTCTTCCGATCTttttcagctcctcctgaacc | 4C-seq *Klf14* distal | This study |
| Klf14-19 F1-P7 2.5 | CAAGCAGAAGACGGCATACGAGATCACTGTGTGACTGGAGTTCAGACGTGTGCTCTTCCGATCTttttcagctcctcctgaacc | 4C-seq *Klf14* distal | This study |
| Klf14-19 F1-P7 2.6 | CAAGCAGAAGACGGCATACGAGATATTGGCGTGACTGGAGTTCAGACGTGTGCTCTTCCGATCTttttcagctcctcctgaacc | 4C-seq *Klf14* distal | This study |
| Klf14-19 F1-P7 2.7 | CAAGCAGAAGACGGCATACGAGATGATCTGGTGACTGGAGTTCAGACGTGTGCTCTTCCGATCTttttcagctcctcctgaacc | 4C-seq *Klf14* distal | This study |
| Klf14-19 R1-P5 | AATGATACGGCGACCACCGAGATCTACACTCTTTCCCTACACGACGCTCTTCCGATCTtcttcaccaaaataccactagaatg | 4C-seq *Klf14* distal | This study |
| **4C-gDMR viewpoint** | | | |
| 4CSeq_Mest_F1 | AATGATACGGCGACCACCGAACACTCTTTCCCTACACGACGCTCTTCCGATCTctggttttcctccagaagga | 4C-seq *Mest* gDMR | This study |
| 4CSeq_Mest_R1 | CAAGCAGAAGACGGCATACGAaggtgtgtcagtgaggag | 4C-seq *Mest* gDMR | This study |
